# YAP1 Depletion Enhances TAZ and its Complexation with TEAD4 and AP-1 Heterodimer C-JUN/FOSB to Promote Gastric Cancer Progression and Metastases

**DOI:** 10.1101/2025.11.13.683933

**Authors:** Jingjing Wu, Dipti Athavale, Curt Balch, Junsong Zhao, Gengyi Zou, Yibo Fan, Yanting Zhang, Joseph Zhao, Mikel Ghelfi, Anthony Pompetti, Gennaro Calendo, Ailing Scott, Shan Shao, Xiaodan Yao, Melissa Pool Pizzi, Christopher Vellano, Vladimir Khazak, Sheng Zhang, Timothy A Yap, Shilpa S Dhar, Raghav Sundar, Francis Spitz, Generosa Graana, Jaffer A. Ajani, Shumei Song

**Affiliations:** Department of GI Medical Oncology, Houston, TX 77030, USA; Coriell Institute for Medical Research, 403 Haddon Ave, Camden, NJ, 08103, USA; Camden Cancer Research Center, 403 Haddon Ave, Camden, NJ, 08103, USA; Gastroesophageal Cancer Program, Yale School of Medicine and Yale Cancer Center, PO Box 208028 New Haven, CT 06520-8028; Departments of Biomedical Sciences and Surgery, Cooper Medical School of Rowan University, 401 Broadway, Camden, NJ, 08103, USA; Department of Pathology, The First Affiliated Hospital of Fujian Medical University, Fuzhou, Fujian, China, 350005; Research Planning and Dev Traction, Houston, TX 77030, USA; NexusPharma Inc. 17 Black Forest Road, Hamilton, NJ 08691; Department of Investigational Cancer (Phase 1 program) at The University of Texas MD Anderson Cancer Center, Houston, TX 77030, USA; MD Anderson Cancer Center at Cooper, Cooper University Hospital, 2 Cooper Plaza, Camden, NJ, 08103, USA

**Keywords:** antisense oligonucleotide (ASO), gastric cancer peritoneal metastasis (GCPM), Hippo pathway, YAP1, TAZ, TEADs1-4, targeted therapy

## Abstract

**Background:** Dysregulation of the Hippo signaling pathway, characterized by aberrant activation of the transcriptional coactivators YAP1 and TAZ, drives tumour progression, immunosuppression and metastasis. Hippo pathway components are emerging therapeutic targets in several solid tumours, however, the expression profiles of Hippo coactivators YAP1, TAZ and their transcriptional factors TEAD1-4 in gastric cancer peritoneal metastases (GCPMs) and their therapeutic value are unknown.

**Objective:** To determine expression status of YAP1, TAZ and TEAD1-4 in GCPMs; and to evaluate whether dual targeting of YAP1 and TAZ provides superior antitumour activity compared with inhibition of either coactivator alone.

**Design:** Expression of YAP1, TAZ and TEAD1–4 was examined in GCPMs by single-cell RNA sequencing and co-immunofluorescent staining. Functional studies using genetic knockout and antisense oligonucleotide (ASO) inhibition of YAP1 or TAZ were performed to assess antineoplastic effects in vitro and in vivo. Co-immunoprecipitation and luciferase reporter assays were used to characterize YAP1/TAZ interactions with TEADs and AP-1 components (JUN and FOSB) and to quantify transcriptional activity. Antitumour efficacy was validated in patient-derived xenograft (PDX) and KP-Luc2 syngeneic models.

**Results:** YAP1, TAZ, and TEADs1- 4 were highly coexpressed in GCPMs and correlated with poor survival. YAP1 inhibition alone elicited compensatory upregulation of TAZ, while combined inhibition of both coactivators maximally repressed cell proliferation and invasion *in vitro*, and tumor growth *in vivo*. Increased TAZ complexation with TEAD4 and AP-1 (c-JUN and FOSB) heterodimer was observed following YAP1 knockdown or pharmacological ASO inhibition. Dual inhibition of YAP1 and TAZ was required to maximally suppress YAP1/TAZ expression and reduce their nuclear accumulation, transactivation of TEAD, and activation of downstream genes.

**Conclusions:** These findings show that combined YAP1 and TAZ inhibition holds promise for the treatment of GCPMs, a highly lethal disease with an urgent need for novel treatment options.

**WHAT IS ALREADY KNOWN ON THIS TOPIC:**

- Gastric cancer with peritoneal metastasis (GCPM), occurring in > 45% of gastric cancer (GC) patients, is a highly lethal malignancy with limited therapeutic options.
- The Hippo pathway mediator Yes-associated protein 1 (YAP1) is intimately involved in chemoresistance, cancer stemness properties, and the epithelial-to-mesenchymal transition in gastric and other cancers.
- While YAP1 represents a promising therapeutic target, clinical trial of YAP1 antisense oligonucleotides has been disappointing.

**WHAT THIS STUDY ADDS:**

- Hippo coactivators YAP1, TAZ and their main transcription factors TEAD1-4 are markedly upregulated in GCPMs and associated with poor prognosis.
- Antisense targeting of YAP1 results in compensatory upregulation of its paralog TAZ.
- Upon YAP1 depletion, TAZ forms transcriptional complexes with TEAD4 and the AP-1 heterodimer (JUN and FOSB).
- Dual targeting of YAP1 and TAZ, by oligonucleotides, achieves maximal suppression of tumour growth *in vitro* and *in vivo*.

**HOW THIS STUDY MIGHT AFFECT RESEARCH, PRACTICE OR POLICY:**

- These findings provide a strong mechanistic rationale for co-targeting YAP1 and TAZ as a therapeutic approach in metastatic gastric cancer.
- Dual Hippo coactivator inhibition could inform the design of future clinical trials aimed at improving outcomes for patients with GCPMs.

## Introduction

Gastric cancer (GC) is the fifth-leading cause of cancer, worldwide, and the fourth-leading cause of cancer death.^1^ In the U.S. alone, the American Cancer Society estimates 30,300 new gastric cancer cases in 2025, with 10,780 deaths^2,3^. Primary causes of GC include *Helicobacter Pylori* infection, male gender, advanced age, consumption of smoked salted meats, and tobacco use.^4^ About 45% of GC patients present with peritoneal metastases (PM, malignant ascites or implants in the peritoneal cavity), having a five-year survival rate of merely 6.5 months.^1^ In particular, our group found that GC with PM (GCPM) predominantly features Hippo/YAP1 involvement, and that inhibiting YAP1 was strongly antineoplastic and suppressive of GCPM.^5^

The complexity of cancer involves intricate signaling pathways that regulate cell growth, proliferation, cell fate decisions, and tissue homeostasis. Among these pathways is the highly conserved Hippo pathway, a critical regulator of organ size, tissue regeneration, stem cell self-renewal, and tumorigenesis through its co-activators Yes-associated protein 1 (YAP1) and transcriptional co-activator with PDZ-binding motif (TAZ) that interact with the transcription factors TEAD1-TEAD4.^6^ YAP1 and TAZ are encoded by paralogous genes, with 46% amino acid identity.^7^

Central to Hippo function is a series of kinases and transcriptional co-activators that coordinate cellular responses to various cues, including cell-cell contact, mechano-transduction, and soluble factors. Under homeostatic conditions, YAP1 and TAZ are phosphorylated by the Hippo kinase LATS 1/2 (large tumor suppressor kinases) and sequestered in the cytoplasm, by 14-3-3-σ proteins, or degraded.^7^ Since YAP/TAZ are mitogenic effectors, cytoplasmic sequestration/degradation effectively constrains cell growth. Under dysregulated conditions (as in various cancers), however, phosphorylation (and thus, Hippo signaling) is disrupted, resulting in YAP or TAZ nuclear translocation and interaction with TEA domain (TEAD) transcription factors TEAD1-TEAD4, activating genes involved in stemness, proliferation, epithelial-to-mesenchymal transition (EMT), angiogenesis, and therapy-resistance.^8^ Adding additional complexity, YAP1 and TAZ can crosstalk with other signaling pathways such as Wnt/β-catenin, AP-1, Notch, STAT3, and TGF-β, amplifying their oncogenic potential and ability to form a permissive microenvironment for tumor progression^7,9–13^.

While YAP1 and TAZ are believed to have considerable functional redundancy, the degree of overlap, in association with cancer phenotypes, remains unclear^7,9,14,12,13^. For example, in hepatocellular cancer (HCC) cells, TAZ knockdown resulted in increased YAP1 levels, concomitant with increased cancer stem cell traits (therapy-resistance and tumorigenicity), and combined YAP1/TAZ inhibition was found most efficacious in tumor suppression.^15^ Likewise, in embryonic development, only combined inhibition of YAP and TAZ could completely halt blastocyst formation, compared to either alone.^16^ Thus, while most Hippo-targeting strategies have focused on inhibition of YAP1, it is possible that this might be insufficient, due to potential compensatory signaling by TAZ. A recent clinical trial (NCT04659096) targeting YAP1 alone using a YAP1 antisense oligonucleotide (ASO)^17^ in advanced solid tumors has failed. However, the reasons for this trial failure and the complementary mechanism for activation of TAZ and its binding partners in GCPM are unknown.

In this study, we extensively characterized co-expression of YAP1 and TAZ in advanced GCPM samples; both interacted with TEAD1-TEAD4 to enhance TEAD transcriptional activity. Further, we found that YAP1 or TAZ ASOs efficiently knocked down YAP1 or TAZ respectively *in vitro*. Moreover, we show that genetic knockout (KO) or pharmacological inhibition of YAP1 enhanced TAZ expression and activity while increasing its interaction with TEAD4 and the AP-1 heterodimer (c-JUN/FOSB). Consequently, combined ASO inhibition of YAP1 and TAZ was necessary to maximally suppress both invasion and proliferation *in vitro* and suppress tumor growth *in vivo*. Thus, combined inhibition of both Hippo signal effectors might be advantageous, over suppression of either alone.

## MATERIAL AND METHODS

The detail materials and methods can be found from online supplemental materials and methods.

## RESULTS

### YAP and TAZ are highly coexpressed in primary and metastatic GCs

As YAP1 and TAZ have previously been reported to cooperate as transcriptional coactivators in malignancies,^6^ we examined their expression in GC tumors, compared to normal tissues. As shown in Figure 1A, mRNAs for both genes were elevated in GC tumors compared to normal tissues. Interestingly, both genes significantly positively correlated (R=0.42; P value=0, Figure 1B) in primary GC tumor tissues (gepia.cancer-pku.cn). Increased expression of YAP1 or TAZ in GC tumor tissues compared to normal was further validated in two independent GC dataset cohorts (GES33335 and GSE146996) (Figure 1C; Supplemental Figure 1A); and both YAP1 and TAZ are highly correlated in primary GC tissues (Figure 1D; Supplemental Figure 1B). The significant correlation of YAP1 and TAZ expression in primary GC tissues was further confirmed in our own TMA (Figures 1E and Supplemental Figure 1C) and in another independent GC cohort (GSE237876; Figure 1F). Further, in our established 18 PDXs from gastroesophageal cancers, both YAP1 and TAZ are highly expressed in these PDXs by protein level (Figure 1G and Supplemental Figures 1D) and mRNA level (Supplemental Figure 1E).

**Figure 1.**
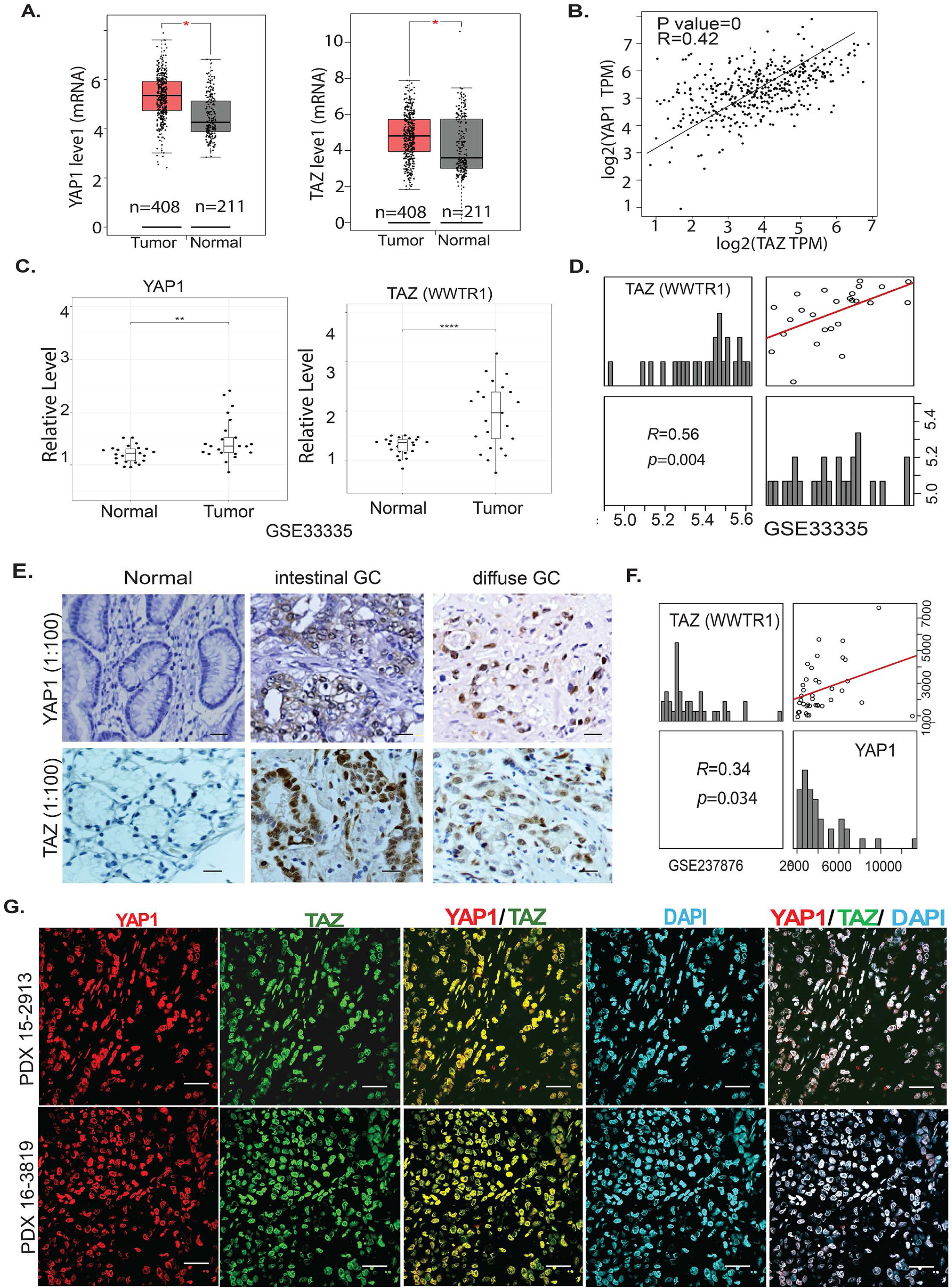
YAP1 and TAZ are highly co-expressed and correlated in primary GC tissues and PDXs. **A**. Increased YAP1 or TAZ in GC tumor tissues compared to normal tissues (http://gepia.cancer-pku.cn). **B**. Both YAP1 and TAZ are significantly correlated (R=0.42; *P* value=0). **C**&**D**. Expression of YAP1 or TAZ is significantly higher in GC tumor tissues than normal and significantly positively correlated (R=0.56, *P*=0.004) in GSE33335 cohort (\*\**P*<0.01; \*\*\*\**P*<0.001); **E**. Expression of YAP1 or TAZ was determined using immunohistochemistry in 390 cases of GC specimen in both intestinal GC and diffuse GC. **F**. The correlation of YAP1 and TAZ in GC tissues was further confirmed in an individual GC cohort (GSE237876) (R=0.34, *P*=0.034). G. co-localization of YAP1 and TAZ in two representative PDXs GC tissues was determined by co-immunofluorescent staining of YAP1, TAZ and DAPI (scale bar: 25μm).

To determine the expression of YAP1 or TAZ and their correlation in GC peritoneal metastases (GCPM), dot plots of scRNASeq from 20 GCPM samples revealed that both YAP1 and TAZ are highly expressed and correlated in tumor cell clusters (Figure 2A). Among 13 tumor cell clusters, YAP1 and TAZ were highly co-expressed and correlated using U Map plots (Figure 2B). Further, we co-stained YAP1 and TAZ in more than 110 GCPM cases using co-immunofluorescent staining (Co-IF) and revealed co-expression of YAP1 and TAZ in most malignant ascites of GCPM samples. As shown by representative staining of both YAP and TAZ in GCPMs (Figure 2C and Supplemental Figure 2A), we noticed that both YAP1 and TAZ highly co-localized in the nuclei of malignant ascites tumor cells in GCPMs. Expression of YAP1 and TAZ in tumor cells than other cell types were validated using co-IF staining with tumor marker EpCAM, stromal marker vimentin or macrophage marker CD163 (Supplemental Figure 2B). Furthermore, two independent GC cohorts with peritoneal metastases further confirmed the much higher positive correlation between YAP1 and TAZ (*WWTR1*) in metastatic lesions of GC (GES237876, R=0.70; GSE289037, R=0.89) (Figure 2D) than that in primary GC cohorts (Figure 1D, Figure 1F and Supplemental Figure 1B); and YAP1 is significantly higher in GC metastasis than primary tumors (Figure 2E). Similarly, TAZ expression significantly increased in GC with peritoneal metastases compared to GC without peritoneal metastasis in an independent cohort (Supplemental Figure 2C). All these data suggest that both YAP1 and TAZ coexist in GC and highly associated with GCPMs.

**Figure 2.**
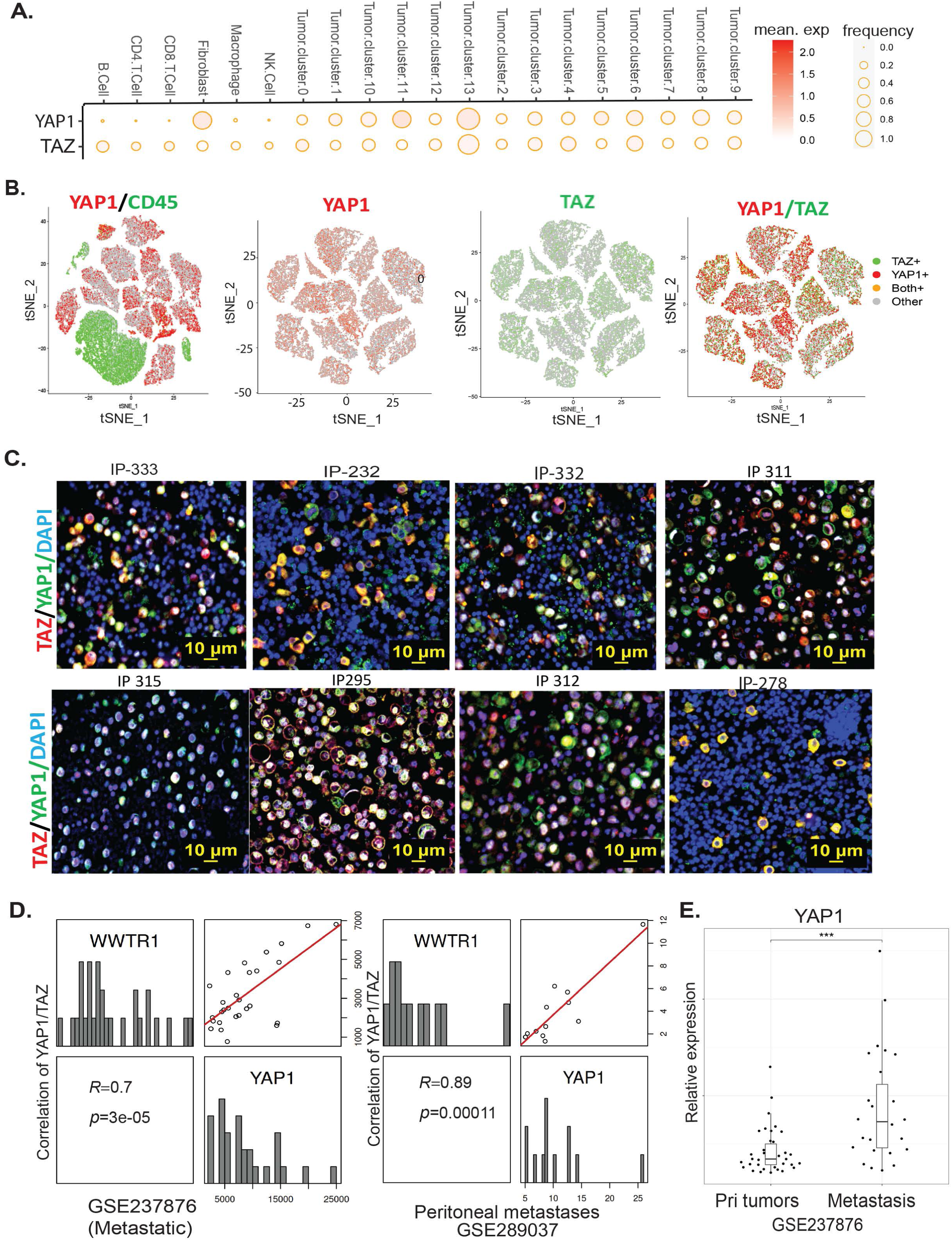
Both YAP1 and TAZ are highly expressed and correlated in metastatic GCPM cells/tissues. **A**. expression of YAP1 and TAZ in tumor cell clusters was shown in dotted plots in 20 GCPM samples by scRNAseq. B. Coexpression of YAP1 and TAZ in 13 tumor cell clusters was shown in tSNE plots in GCPM samples by scRNAseq **C**. Expression of YAP1 and TAZ as determined by dual-immunofluorescent staining (co-IF) in representative GCPM samples. Scale bar: 10μm. **D**. High correlation of YAP1 and TAZ in two metastatic GC cohorts (GSE237876 and GSE289037). **E.** Expression of YAP1 was significantly increased in peritoneal metastasis compared to primary tumors from a GC cohort (GSE237876).

### YAP and TAZ complex with TEAD1-TEAD4 and synergically activate TEAD transcriptional activity

It is well-established that during developmental processes, YAP1 and TAZ, which lack DNA-binding capacity, are activated by loss of phosphorylation, translocate to the nucleus, and transactivate target growth genes through their TEAD transcription binding partners. Four TEAD members including TEAD1, TEAD2, TEAD3 and TEAD4 can be utilized by coactivators YAP1 or TAZ or both. However, expression of these TEADs in GCPM and their association with YAP1 or TAZ, and their clinical relevance are unknown. First, RNAseq analysis in 20 GCPM samples revealed that four TEADs are highly enriched in 13 tumor cell clusters with TEAD4 enriched in tumor cluster 12 that highly associated with poor survival, while TEAD1 and TEAD2 were also expressed highly in cancer-associated fibroblasts (Figure 3A). Further, we revealed that TEAD1-4 are highly increased in GC tumor tissues compared to normal from 2 independent GC cohorts GES33335 and GES146996 (Supplemental Figures 3A&3B). To explore the four TEADs clinical relevance, we explored the TCGA dataset and found that elevated expression of all four TEADs significantly associated with short GC patient survival respectively in GC samples that contain more than 600 cases (Figure 3B).

**Figure 3.**
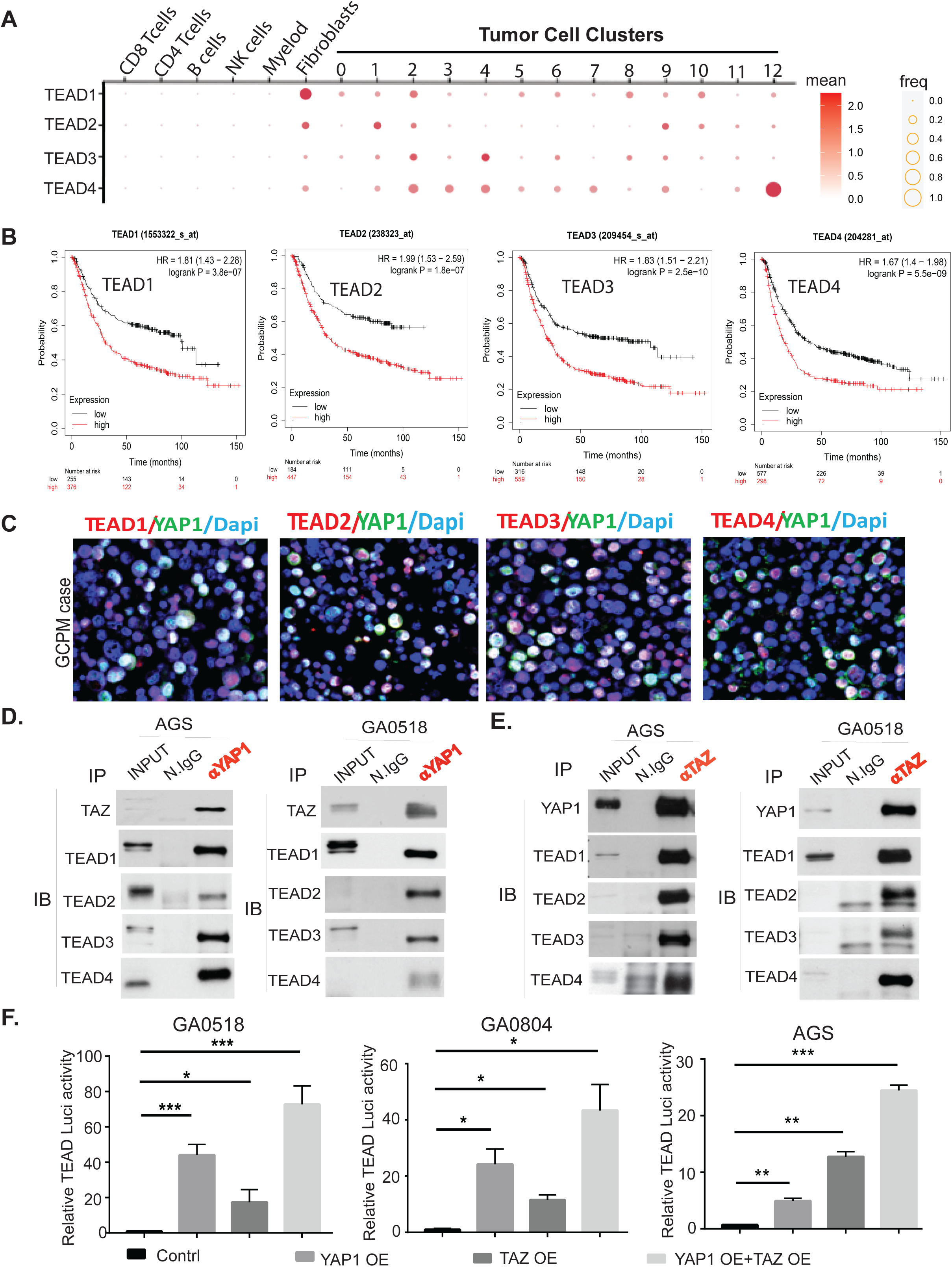
High expression of TEAD1-TEAD4, the transcription factors (TFs) for YAP1/TAZ in GCPM associates with poor survival; **A**. scRNAseq showed the expression of TEAD1, TEAD2, TEAD3 and TEAD4 by dot plots from 20 GCPM specimens; **B**. Association of TEAD1-TEAD4 expression with GC patients’ survival in more than 600 advanced GC patients respectively from the TCGA database (kmplot.com). **C**. Representative GCPM cases stained by dual-immunofluorescence of YAP1 with TEAD1-TEAD4 respectively. **D**. YAP1 interacts with TAZ and TEAD1-TEAD4 in AGS and GA0518 GC cells as shown by co-immunoprecipitation pulldown using anti-YAP1 antibody. **E**. TAZ interacts with YAP1 and TEAD1-TEAD4 as shown by co-immunoprecipitation pulldown using anti-TAZ antibody in AGS and GA0518 GC cells. **F**. TEAD transcriptional activity was determined as previously^18^ by transient cotransfection of 5X UAS-luciferase reporter and Gal4-TEAD4 with either YAP1cDNA or TAZ cDNA or both in GA0518, GA0804 and AGS cells for 48 hours. *p<0.05, **p<0.01; ***p<0.001.

To further detect if TEAD is actively expressed in GCPM samples, we co-stained TEAD1-TEAD4 and YAP1 using co-immunofluorescent staining (co-IF) in our GCPM specimens and found that four TEADs including TEAD1-TEAD4 are highly expressed and colocalized with YAP1 (Figure 3C). To elucidate the interaction of these TEADs with their co-activator YAP or TAZ, we performed co-immunoprecipitation using either specific YAP1 antibody or TAZ antibody. As shown in Figure 3D, an anti-YAP1 antibody coimmunoprecipitated TAZ and TEAD1-TEAD4, with reciprocation shown by anti-TAZ antibody pulldown of YAP1 and TEAD1-TEAD4 (Figure 3E) indicating that YAP1 actively interacts with TAZ and both complex with TEAD1-TEAD4 in two GC cell lines. Functional cooperation of YAP1 and TAZ in GCPMs was shown in transactivation, with combined YAP1 and TAZ overexpression (OE) maximally driving TEAD luciferase activity, compared to OE of YAP1 or TAZ alone in two GCPM derived tumor cells (GA0518 and GA0804) and AGS GC cell line (Figure 2F). Co-localization of YAP1 or TAZ with TEADs in representative GCPM samples was further confirmed by co-IF (Supplemental Figure 3C). Altogether, these data suggest that YAP1, TAZ and TEAD1-TEAD4 are highly expressed and associated in GCPM samples; thus, the YAP1/TAZ/TEADs axis could be a promising therapeutic target in GCPM.

### YAP1 or TAZ antisense oligos (ASO) effectively suppress their expression and potently inhibit invasion of patient-derived GCPM cells

We previously showed that in gastroesophageal cancer, inhibition of YAP1 was antitumorigenic, using genetic CRISPR/CAS9 knockout or a pharmacologic inhibitor^5,18^. However, ASOs have the advantages of depletion of the actual protein of interest, minimal off-target effects, and stabilization by chemical modifications, and have now been under preclinical and clinical examination and optimization for over two decades^19,20–22^. Thus, we examined three distinct ASOs for knockdown of YAP1; ASO#3 was the most efficacious for suppressing YAP1 expression and subsequently inhibiting transactivation of the YAP1 target gene *Cyr61* in both GA0518 and GA084 patient-derived GCPM cells (Figures 4A-4D) and also dramatically reduced SOX9 and BIRC5, the two reported YAP/TEAD targets in a dose dependent manner (Supplemental Figure 4). Correspondingly, YAP1 ASO significantly blocked tumor cell invasion in both GA0518 and GA0804 in a dose-dependent manner (Figure 4E). Consistent with YAP1’s oncogenic functions, YAP1 ASO effectively inhibited tumor cell proliferation in GA0518cells, while YAP1 KO clones failed to respond to YAP1 ASO indicating that YAP1 ASO antitumor effects rely on YAP1 expression (Figure 4F). In testing additional YAP1 ASO#10 and YAP1 ASO#11, we found that either YAP1 ASO#10 or YAP1 ASO#11 dramatically reduced colony formation in a dose dependent manner in GA0804 cells (Supplemental Figure 4B).

**Figure 4.**
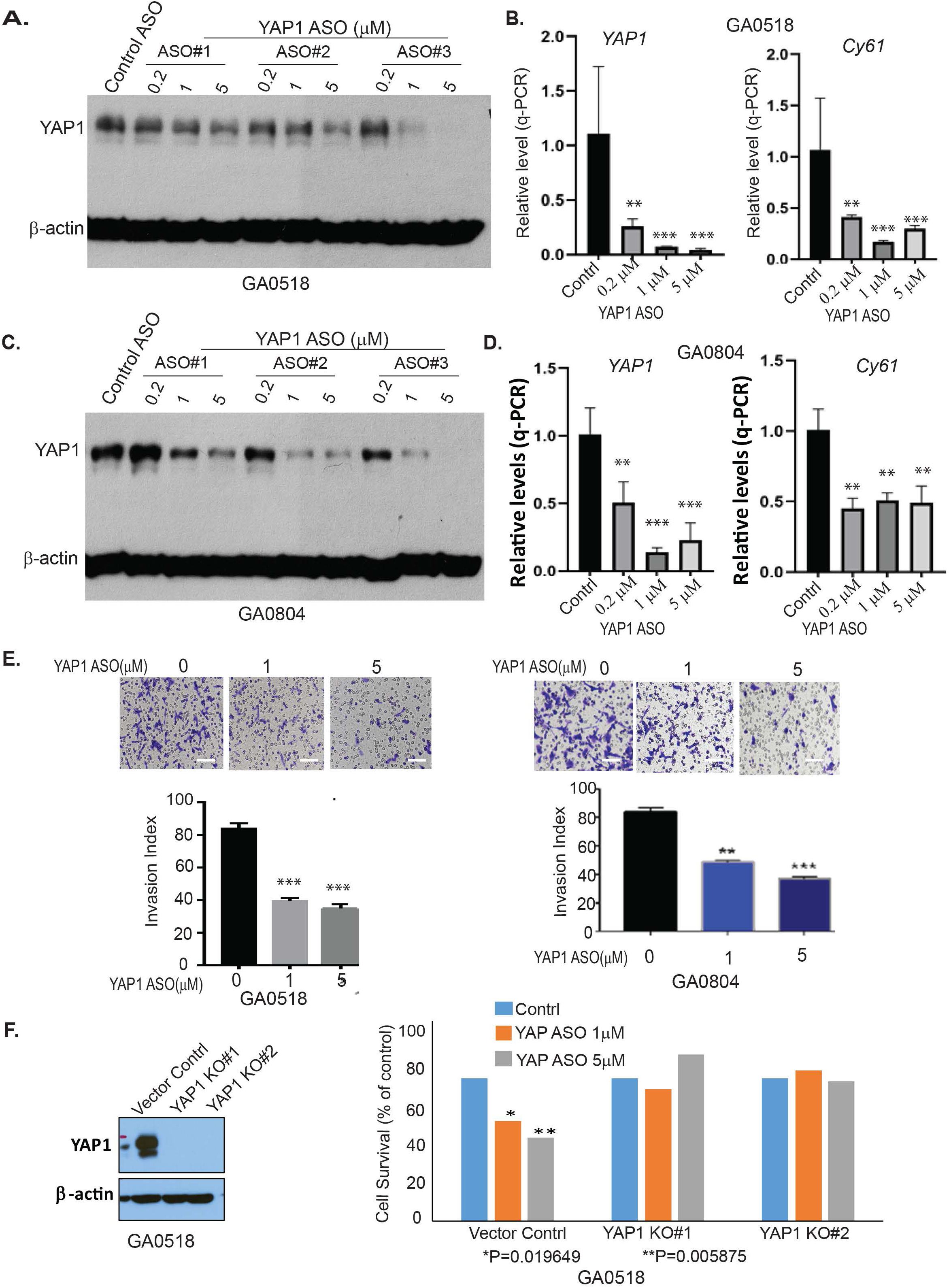
Antisense oligonucleotides (ASO) of YAP specifically suppress YAP expression, transcription, and reduce tumor cell malignant behaviors *in vitro*. A.YAP1 protein was determined by Western blot after treatment with YAP1 ASOs in GA0518 cells as indicated; **B.** YAP1 mRNA (left) and its target Cyr61 (right) mRNA levels were dose-dependently decreased by YAP1 ASO as determined by q-PCR in GA0518 cells **p<0.01; ***p<0.001; **C**. YAP1 protein was determined by Western blot after treatment with YAP1 ASOs in GA0804 cells; **D.** YAP1 mRNA (left) and its target Cyr61 (right) mRNA levels were decreased by YAP1 ASO as determined by q-PCR in GA0804 cells; **p<0.01; ***p<0.001; **E**. YAP1 ASO significantly inhibited tumor cell invasion in both GA0518 (left) and GA0804 (right) GCPM tumor cells. Lower panels, quantification. **p<0.01; ***p<0.001; **F**. Western blot showing YAP1 knockout efficiency (left); cell survival was detected by MTS assay in GA0518 cells with or without YAP1 KO (right) treated with YAP1 ASO at different dosage. *p<0.05; **p<0.01;

Analogously, we assessed the effects of TAZ ASO on TAZ expression. TAZ ASO showed dose-dependent downregulation of TAZ protein (Figure 5A) and mRNA (Figure 5B) in both GA0518 and GA0804 patient-derived tumor cells. The specificity of TAZ ASO in decreasing TAZ but not YAP1 protein is shown in Supplemental Figure 5A using a dual antibody recognizing both YAP1 and TAZ. Furthermore, we noticed that TAZ but not YAP1 ASOs significantly downregulated TEAD luciferase activity induced by co-transfection of TAZ and TEAD indicating that TAZ ASO specifically inhibits TAZ-induced TEAD activation in GA0518 and AGS cells (Figure 5C), while both YAP1 ASO or TAZ ASO significantly reduced YAP- and TAZ-coactivated TEAD transcriptional activity following co-overexpression of YAP1, TAZ and TEAD with a UAS luciferase plasmid (Figure 5D). In contrast, overexpression of TAZ dramatically increased TEAD transcriptional activity in GA0518 cells (Figure 5E). Correspondingly, TAZ ASO dramatically suppressed GC tumor cell invasion in both GA0518 and GA0804 cells (Figure 5F). One TAZ ASO (#896558) dramatically reduced an aggressive GA0518 GCPM cells’ colony formation as comparable to CA3, a reported potent YAP/TEAD inhibitor^18^ (Supplemental Figures 5B&5C). All together, these data support the feasibility of antisense inhibition of YAP1 or TAZ expression and their oncogenic function in GCPMs.

**Figure 5.**
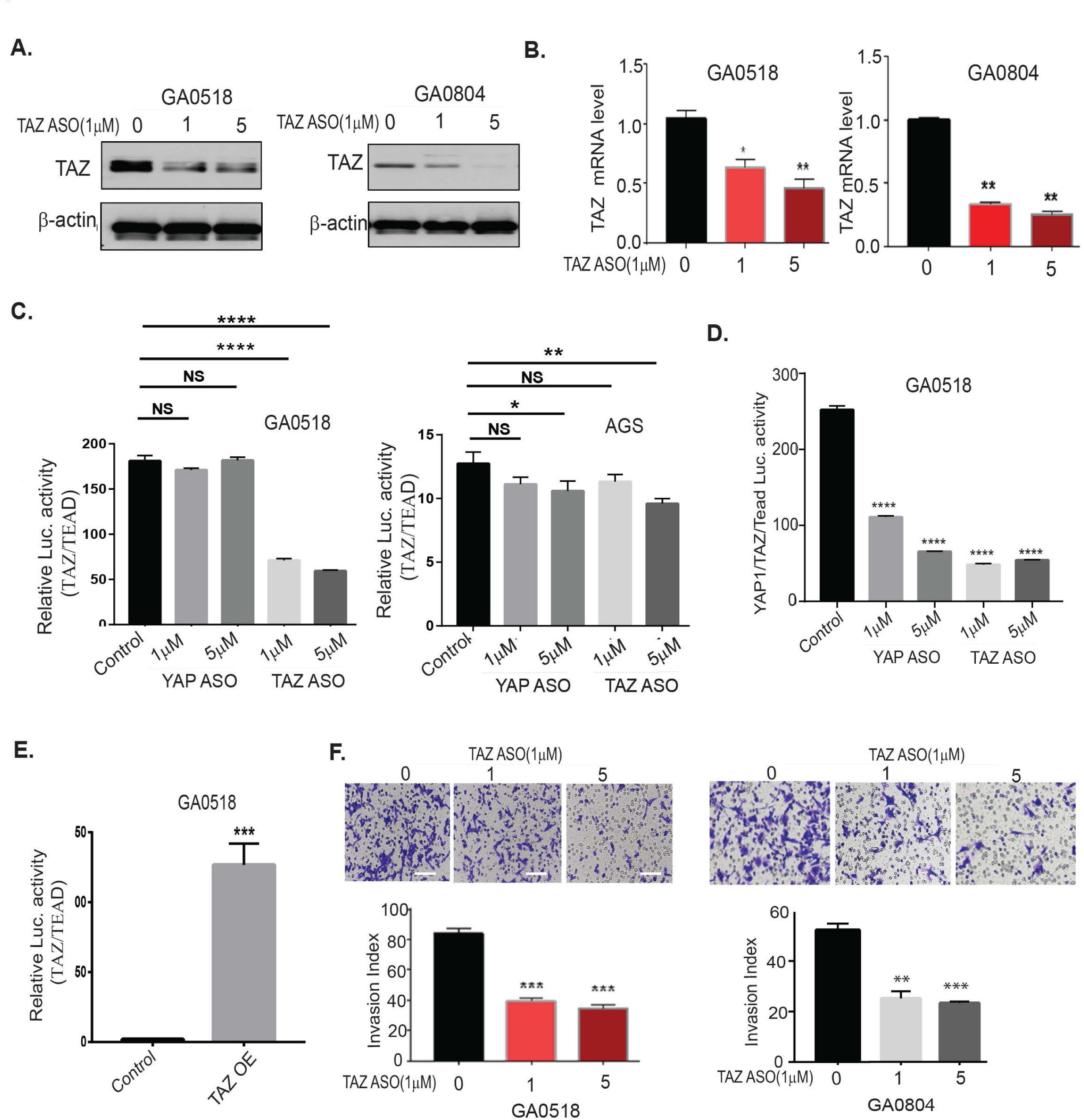
TAZ ASO suppresses TAZ expression, transcription, TAZ/TEAD transcriptional activity and reduces tumor cell invasion *in vitro*. A&B. TAZ protein and mRNA levels were decreased by TAZ ASO as determined by Western blot and q-PCR in both GA0518 and GA0804 cells; *p<0.05, **p<0.01; **C**. TAZ/TEAD transcriptional activity was determined as previously^18^ by transient cotransfection of 5X UAS-luciferase reporter and Gal4-TEAD4 with either TAZ cDNA or YAP1cDNA in GA0518 and GA0804 for 48 hours. *p<0.05, **p<0.01; ****p<0.001. **D**. Luciferase activity of YAP1/TAZ/TEAD was determined after cotransfection of 5X UAS-luciferase reporter and Gal4-TEAD4 with both TAZ cDNA and YAP1cDNA and then treated with YAP1 ASO or TAZ ASO in GA0518 cells; ****p<0.001; **E**. Luciferase activity of TAZ/TEAD was determined after cotransfection of 5X UAS-luciferase reporter and Gal4-TEAD4 with TAZ cDNA overexpression as done previously. ***p<0.001; **F**. TAZ ASO strongly suppressed tumor cell invasion capacity in a dose-dependent manner in both GA0518 (left) and GA0804 (right) cells. **p<0.01; ***p<0.001.

### YAP1 depletion upregulates TAZ and enhances TAZ interactions with TEAD4 and the AP-1 heterodimer of C-JUN/FOSB

While transcriptional regulation of YAP1 and TAZ is complex, involving crosstalk with numerous pathways, we and others^15^ have shown that in addition to binding TEAD transcription factors, YAP1/TAZ can interact with other transcription factors/mediators such as STAT3, β-Catenin, Notch, BRD4, SMAD3 and AP-1 to mediate their oncogenic functions.^9,12–14,23,24^. Consequently, inhibition of TEAD alone may be insufficient, as YAP1 or TAZ can engage alternative transcription factors to drive target gene expression, even though current therapeutic approaches targeting the Hippo/YAP1/TEAD axis remain focused on TEAD inhibition^25–27^. However, how YAP1 inhibition influences TAZ interaction with its binding partners in GCPMs is unclear. One could hypothesize that depletion of YAP1 could be compensated by upregulation of TAZ and increased TAZ binding to other TFs in addition to TEAD, and vice versa. As shown in Figure 5A, genetic YAP1 knock out (KO) upregulated TAZ protein and mRNA in GA0518 cells (Figure 6A). Similarly, ASO inhibition of YAP1 increased TAZ protein and mRNA levels in both GA0518 and GA0804 cells (Figure 6B). Interestingly, using co-immunoprecipitation, we found that YAP1 genetic KO or YAP1 ASO treatment in GA5018 cells dramatically increased TAZ complexation with TEAD4 and the AP-1 heterodimer (c-JUN and FOSB) (Figure 6C&6D). The co-localization of TAZ and TEAD4 was further validated in our representative GCPMs specimen (Figure 6E). Moreover, upregulation of TEAD4 and AP-1 heterodimers c-JUN and FOS-B upon YAP1 depletion in GA0518 cells was further validated using co-IF (Figure 6F). These results indicate that YAP1 depletion either by genetic KO or ASO pharmacological inhibition upregulates TAZ and increases its binding to TEAD4 and the AP-1 components c-JUN and FOSB that facilitate GC progression and metastasis. This may explain why targeting YAP1 alone in clinical trials failed thus providing rationale for combination therapeutic strategies against GCPMs.

**Figure 6.**
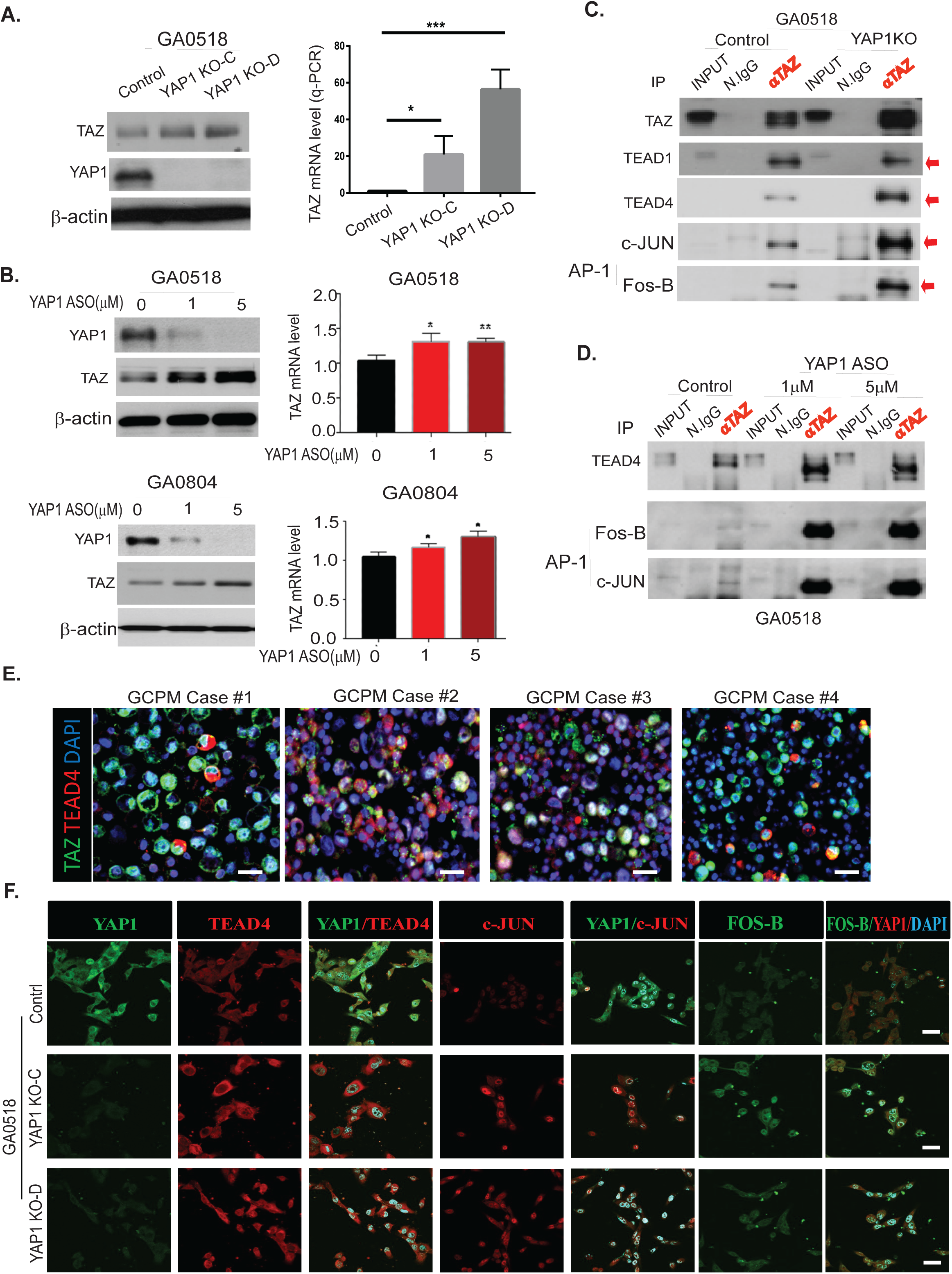
Inhibition of YAP1 by Lenti-CRISPR/CAS9 or YAP1 ASO increases TAZ expression, transcription and increased TAZ binding to TEAD4 and the AP-1 heterodimer C-Jun/FOSB in GC cells. **A.** TAZ level increased upon YAP1 KO by Western blot (left) and q-PCR (right) in GA0518 cells; **B.** YAP1 ASO decreased YAP1 but increased TAZ protein (left) and mRNA (right) levels in both GA0518 (above) and GA0804 (below) cells by Western blot and q-PCR respectively; **C**. YAP1 KO dramatically increased TAZ binding to TEAD4 and the AP1 heterodimer of C-JUN and FOSB as determined by co-IP in GA0518 cells; **D**. Similarly, inhibition of YAP1 by ASO increased TAZ binding to TEAD4 and the AP1 heterodimer of C-JUN and FOSB as determined by co-IP in GA0518 cells. **E**. Co-IF staining of TAZ and TEAD4 in four representative GCPM specimen. Scale bar: 25μm. **F**. Expression of YAP1, TEAD4, c-JUN and FOS-B was determined by co-IF using their specific antibody respectively in GA0518 YAP1 KO clones compared to GA0518 Control cells. Scale bar: 20μm.

### Combined inhibition of YAP1 and TAZ represses their expression and suppresses tumor cell malignant behaviors *in vitro*

Previous studies showed that combined inhibition of YAP1 and TAZ was necessary to prevent blastocyst formation (versus either protein alone), during embryonic development,^16^ and repression of hepatocellular tumorigenesis.^15^ Consequently, we assessed whether this combination could maximally repress each paralog’s expression and function in GC cells. As depicted in Figure 7A-7D, YAP1 ASO treatment alone dose-dependently decreased YAP1 protein and mRNA in GA0518 cells (Figures 7A&7B), while complementarily increasing TAZ protein and mRNA levels as mentioned earlier. Vise versa, ASO inhibition of TAZ alone dramatically decreased TAZ protein and mRNA levels, while increasing YAP expression in both GA0518 and GA0804 cells. However, combined inhibition of YAP1 and TAZ using their ASOs reduced both expression of YAP1 and TAZ in both cell lines in protein and mRNA level (Figures 7A-7D and Supplemental 6A). Correspondingly, combined ASO inhibition of YAP1 and TAZ dramatically reduced tumor cell malignant behaviors including tumor cell invasion and colony formation (Figures 7E&7F) and the combined inhibition of both YAP1 and TAZ elicited the best anti-tumor cell proliferation effects in GA0518 and GA0804 patient-derived tumor cells with YAP1 high expression but less so in the YAP1 none cell line MKN45 cells (Supplemental Figure 6B-6D). These data indicate that co-targeting YAP1 and TAZ is the best strategy to treat GCPMs with hyperactivation of Hippo/YAP1/TAZ signaling.

**Figure 7.**
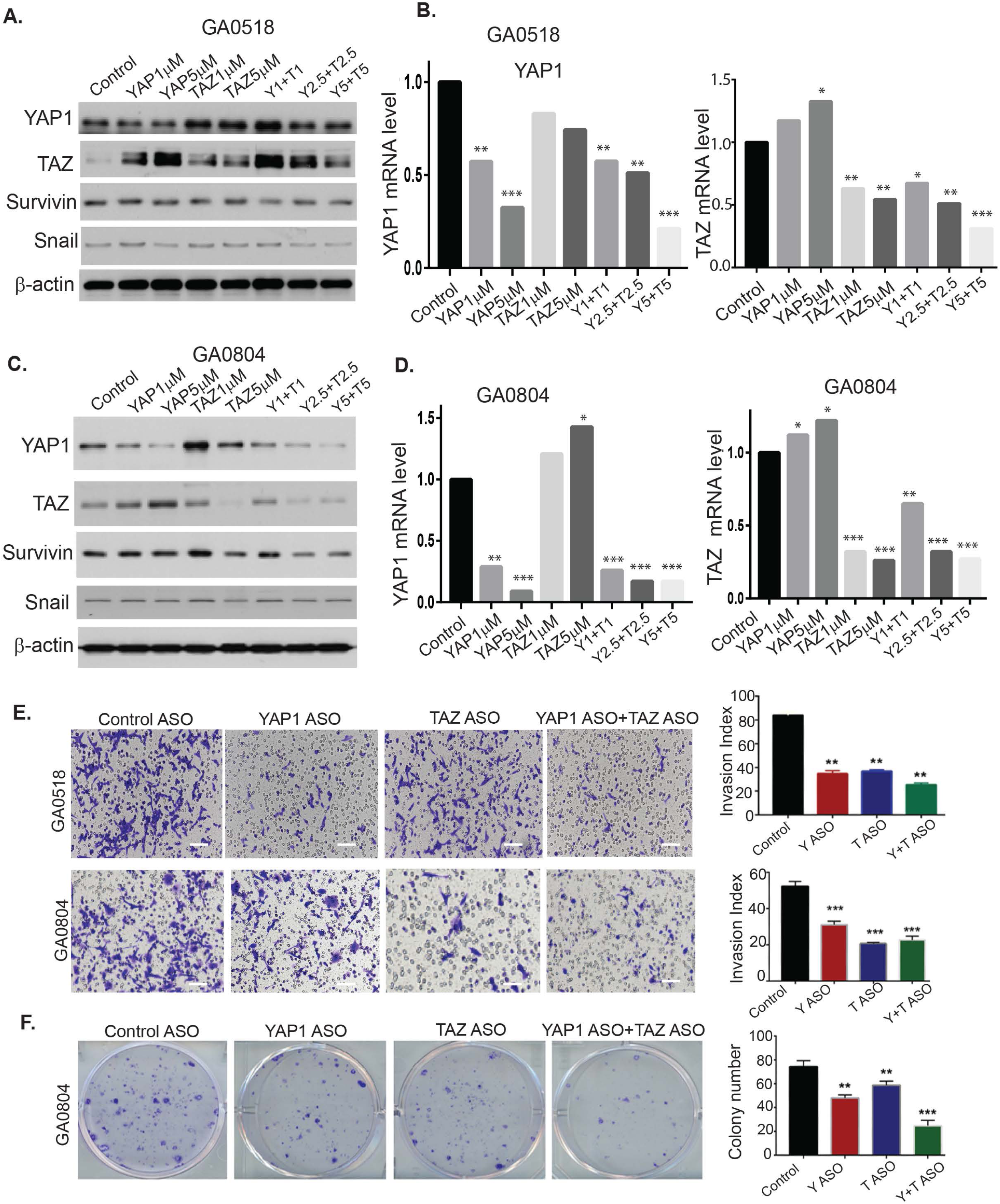
ASO co-targeting of YAP and TAZ dramatically suppresses both YAP1 and TAZ expression and tumor cell invasion and colony formation in patient-derived GCPM cells. **A-D.** YAP1 and TAZ protein and mRNA levels were determined by western blot and q-PCR upon treatment of YAP1 ASO or TAZ ASO or their combination in both GA0518 and GA0804 cells. *p<0.05, **p<0.01; ***p<0.001. **E.** Invasion capacity was determined by western blot and q-PCR upon treatment of YAP1 ASO or TAZ ASO or their combination in both GA0518 and GA0804 cells. **p<0.01; ***p<0.001; Scale bar: 25μm; **F**. Colony formation was determined in GA0804 cells upon treatment of YAP1 ASO or TAZ ASO or their combination. **p<0.01; ***p<0.001.

### Co-targeting YAP1 and TAZ suppresses tumor progression and sensitizes tumors to anti-PD1 immune therapy *in vivo*

To determine the antitumor effects of ASO co-inhibition of YAP1 and TAZ O *in vivo*, we used an established GA0518 PDX line that highly expresses both YAP1 and TAZ (Supplemental Figure 2B). We injected patient-derived GA0518 PDX line cells subcutaneously and treated mice using YAP1 ASO (40mg/kg, 5 times a week), TAZ ASO (40mg/kg, 5 times a week) or their combination (YAP ASO 20 mg/kg + TAZ ASO 20 mg/kg). After three weeks of treatment, tumor weights were significantly reduced by the combined YAP1 ASO and TAZ ASO treatments, compared to either alone or control cells (Figure 8A). Similarly, tumor volumes were significantly reduced in a time-dependent manner, with a YAP1 ASO and TAZ ASO combination providing the greatest inhibition (Figure 8B). Moreover, while the YAP1 or TAZ ASO groups greatly reduced the expression of YAP1 or TAZ respectively and the KI67 proliferation marker, the combination YAP1 and TAZ co-targeting group most suppressed YAP1 and TAZ expression and KI67+ proliferating populations (Figure 8C). Independent similarly designed experiments treating additional PDX, using different YAP1 and TAZ ASOs revealed the consistent results that the combination of YAP1 ASO and TAZ ASO demonstrated the best anti-tumor effects than either treatment alone (Supplemental Figures 7A-7B). The expressions of YAP, TAZ, epithelial tumor marker EpCAM and stromal marker vimentin were dramatically reduced the combination treatment (Supplemental Figure 7C).

**Figure 8.**
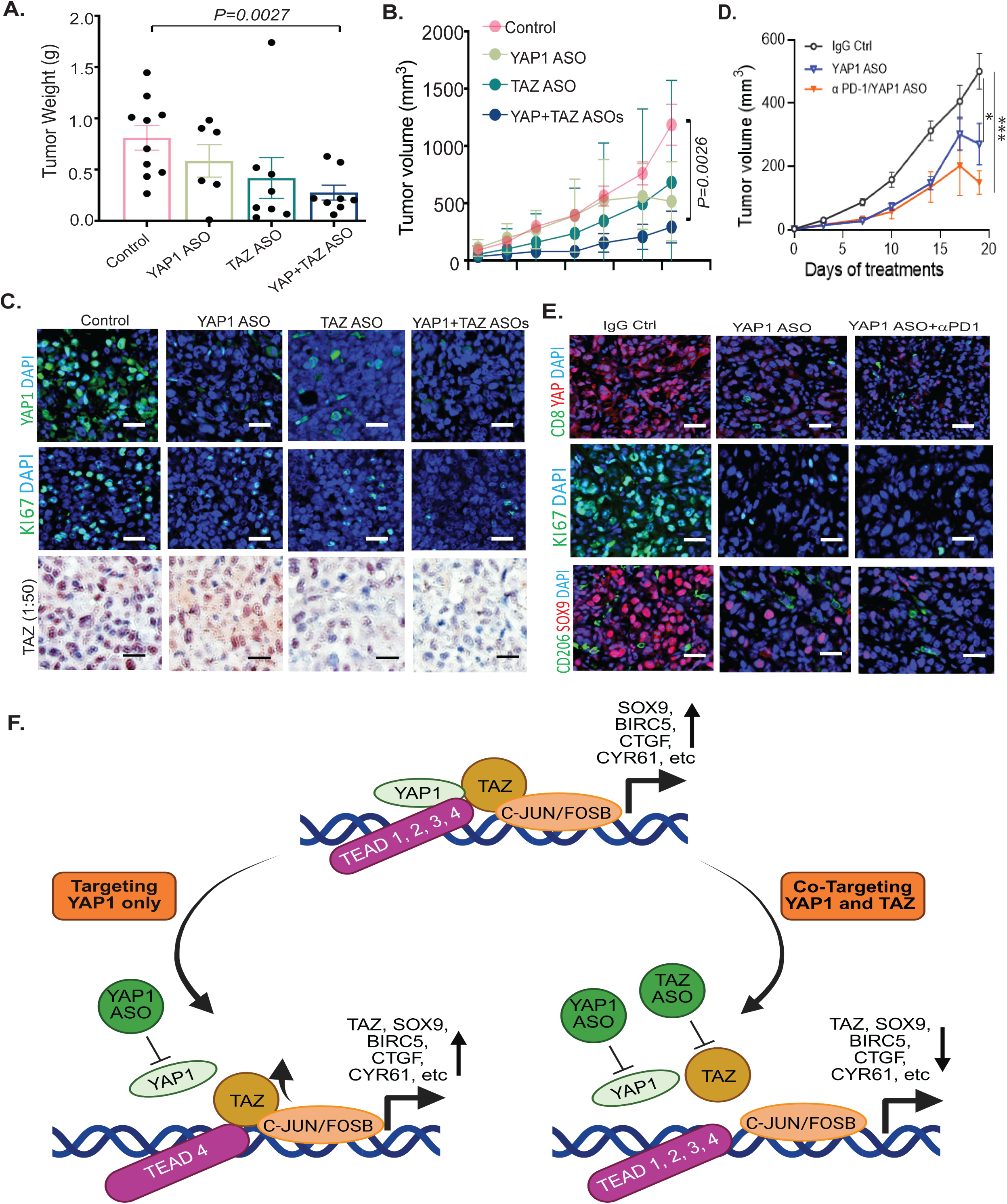
ASO co-targeting of YAP and TAZ significantly attenuated tumor growth *in PDX* with greater antitumor effects when combined with anti-PD1 immunotherapy in the KP-Luc syngeneic model. A-B. Combined ASO inhibition of YAP1 and TAZ suppressed tumor weights and volumes in the GA0518 PDX model; YAP1 ASO or TAZ ASO: 50mg/kg, 5 times a week for three weeks. **C**. Expression of YAP1, TAZ and KI67 as determined by IF staining or immunohistochemistry in YAP1, TAZ, or combination ASO-treated PDX tumors; Scale bar: 20 μm; **D.** Combination of YAP1 ASO and anti-PD1 antibody had greater antitumor effects than YAP1 ASO alone or the control group; *p<0.05, ***p<0.001; **E**. Expressions of YAP1, KI67 and SOX9 were decreased by the YAP1 ASO/anti-PD1 combination treatment, while CD8 infiltration was also increased by the combination treatment over either alone. Scale bar: 20 μm; **F**. Working model and the rationale for co-targeting YAP1 and TAZ in advanced GCPM patients.

To further examine possible immunosensitization, we combined anti-YAP1 ASO with the checkpoint inhibitor anti-PD-1 in KP-Luc2 syngeneic mouse model and found that the combination to be most efficacious in tumor volume inhibition and suppression of the cancer stemness marker SOX9, and proliferation marker Ki67 in addition to strongly suppressing YAP1 expression and increased CD8 T cell infiltration (Figures 8D& 8E). These results show the antitumorigenic effect of combined YAP1/TAZ inhibition, and the possible efficacy of combining anti-YAP1 and anti-PD-1.

## DISCUSSION

Peritoneal metastases (PM; malignant ascites or implants) in GC patients (GCPM) is common and poses a challenge with short survival and lack of effective therapeutics. In this study, using scRNAseq and immunofluorescent staining, we observed that both YAP1, TAZ and their transcriptional factors TEAD1-TEAD4 are highly expressed in GCPM tumor cells; high expression of these proteins associated with poorer prognosis. Further, we note that recently developed YAP or TAZ ASOs can effectively and specifically suppress YAP or TAZ expression and transcription accompanied by decreased tumor cell invasion, colony formation and cell proliferation in YAP1/TAZ high GC cells. Interestingly, ASO inhibition of YAP1 or TAZ alone could complementarily increase the other at the protein and mRNA levels. Moreover, we showed that genetic or pharmacological inhibition of YAP1 enhanced TAZ expression, activity and increased its interaction with TEAD4 and the AP-1 heterodimer (c-JUN/FOSB). Simultaneous ASO inhibition of YAP1 and TAZ reduced both YAP1and TAZ proteins and mRNA levels and their downstream targets while significantly decreasing proliferation and invasive capacity of YAP1 high tumor cells. Most importantly, co-targeting YAP and TAZ by ASOs significantly attenuated tumor progression and GCPM in the PDX model and sensitized tumors to anti-PD1 immunotherapy in the KP-Luc syngeneic model. Taken together, our studies open a new avenue for developing a novel therapeutic strategy of ASO co-targeting of both YAP1 and TAZ in GCPMs.

Dysregulation of the Hippo signaling pathway (with failure to regulate YAP1 and its paralog, TAZ) associates with numerous phenotypes of malignant disease, including the epithelial-to-mesenchymal transition (EMT), angiogenesis, and generation of cancer stem cells.^6^ With 46% identity, YAP and TAZ have been traditionally accepted as interchangeable.^7^ However, a handful of studies have suggested differing, malignancy-specific roles in tumor progression, with TAZ (versus YAP1) preferentially overexpressed in hepatocellular carcinoma (HCC)^15^ and the two proteins playing disparate roles in kidney function.^28^ In another study of gastric cancer cells, using YAP1 or TAZ overexpression combined with knockout of the paralog, although numerous similar target genes/functions were found (including platelet signaling and lipoprotein formation), YAP1 expression was distinctly associated with cell-substrate junctions.^29^ Consequently, one could hypothesize that in many cases, the combined effects of both proteins, versus either alone, would be additive or synergistic. Indeed, it was shown that the combination of YAP1 and TAZ inhibition was necessary to suppress blastocyst formation, during embryonic development,^16^ and for maximal inhibition of tumorigenicity in HCC cells.^15^ Likewise, both YAP1 and TAZ were required for liver regeneration in a genetically engineered mouse model.^30^ Although studies have largely focused on YAP1 function in tumor progression and metastasis, we believe that TAZ is equally important in mediating tumor cell malignant behaviors in GC. However, it is thus far less clear if both YAP1 and TAZ coactivators co-exist in GCPMs and how both cooperate to mediate tumor growth and metastasis. In this study, our findings revealed that both YAP1 and TAZ highly coexist and correlate in GCPM specimens, and overexpression of both YAP1 and TAZ in GC cell lines synergistically increased TEAD transcriptional activity. Combined inhibition of YAP1/TAZ was necessary for maximal downregulation of each, while also most greatly suppressing GC neoplastic phenotypes providing a strong rationale for co-targeting both YAP1 and TAZ coactivators in GCPMs with highly activated YAP1/TAZ/TEAD signaling.

Recently, more attempts have emerged to develop TEAD inhibitors from academia or pharma to combat activation of Hippo/YAP TAZ/TEAD signaling, with several undergoing clinical trials (NCT04665206; NCT05228015). However, when inhibiting TEAD transcriptional factors, the Hippo co-activators YAP1 or TAZ might utilize other transcriptional factors to facilitate tumor cell survival, progression, and metastases. It has been reported that YAP1/TAZ can cooperate with other signaling molecules and bind other TFs such as Wnt/β-catenin, Notch, STAT3, AP-1 and TGF-β/SMAD mediating their oncogenic and metastatic potential^7,9–13^. Furthermore, while most Hippo/YAP/TEAD-targeting strategies have focused on inhibition of YAP1, it is possible that this might be insufficient, due to possible upregulation of TAZ. A recent clinical trial (NCT04659096) targeting YAP1 alone using an YAP1 antisense oligo (ION537)^17^ in advanced solid tumors was terminated due to toxicity and lack of efficacy in advanced solid tumors. Indeed, our findings revealed that TAZ expression and activity increased upon YAP1 inhibition, and TAZ strongly interacted with TEAD4 and the AP-1 heterodimer c-JUN and FOSB to increase their downstream targets that facilitate GC cell growth and metastasis.

A potential interpretation of our findings is depicted in Figure 8F. In this scenario, during tumor progression, YAP1 and TAZ complex with TEAD transcriptional factors (TEAD1-TEAD4) to elicit target gene (e.g., *Cyr61*, *CTGF, Birc5* and *SOX9* etc) expression (top). Inhibition of YAP1 alone using YAP1 ASO upregulates TAZ, possibly via an AP-1 feed forward loop, to maintain tumorigenicity (e.g., proliferation, stemness, EMT, etc.). This could occur via increased TAZ binding to TEAD4 and the AP-1 heterodimer c-JUN and FOSB (lower left), while combined YAP1 and TAZ inhibition reduce both YAP1 and TAZ binding to TEAD and AP-1 TFs to downregulate target genes and cancerous traits (lower right). While others have demonstrated YAP1 compensation following TAZ knockdown,^15^ we believe we are the first to demonstrate the reciprocal effect, i.e., TAZ upregulation following YAP1 suppression. Moreover, while the mechanism of TAZ upregulation, by YAP1 depletion, remains unclear, a feed-forward loop for YAP/TAZ via AP-1 signaling was previously shown in uveal melanoma;^31^ our demonstration of AP-1/TAZ complexation would suggest that this could also positively re-enforce TAZ expression.

In summary, the ancient Hippo pathway, discovered in a screen for tumor inhibitors,^6^ plays an integral role in cancers of the gastrointestinal tract.^32^ While the Hippo effector YAP1 has been intensely investigated as a therapeutic target, expression and activation of its paralog TAZ play critical role to maintain tumorigenic phenotypes especially upon YAP1 depletion, thus diminishing anti-YAP1 efficacy. Further, in targeting TEADs alone which are currently emerging in the preclinical and clinical phases, the Hippo coactivators YAP1 or TAZ might utilize other TFs such as AP-1, BRD4, STAT3 etc. Thus, simultaneously targeting both the YAP1 and TAZ oncoproteins could allow for maximal tumor suppression, therefore warranting further study of such combinations in both preclinical and clinical studies.

## Acknowledgements

This work was supported by grants from the NIH (R01 CA269685), DOD (CA210457 and CA230323) and NJCCR (COCR26RBG006). We thank Ionis Pharmaceuticals, Inc. for kindly providing YAP1 ASOs or TAZ ASOs used in this study.

## Author contributions

All authors are responsible for the overall content as the guarantors and all authors revised and approved the final version of the manuscript. Conception and, design: SS and JAA; Draft the manuscript: SS, CB; development of methodology, acquisition of data (performed experiments, analysed data): JW, DA, JZ, GZ, YF, YZ, JZ, MG, AP, GC, AS, SS, XY, MP, SS; acquired and managed patients, provided facilities, resources, PDXs and other support: CV, VK, SZ, TY, SD, RS, FS, GG, JA, SS; Analysis and interpretation of data: AP, GC, JZ, SS.

## Data availability statement

Data are available in a public, open access repository.

## Patients and Ethics Statement

Ethical approval for this study was obtained from the Institutional Review Board of The University of Texas MD Anderson Cancer Center and the Cooper-Coriell Cancer Biobank (IRB CCCB0001), Camden Cancer Research Center. All patients who volunteered to provide research specimens provided written informed consent using an approved document.

## Materials and Methods

### Cells and Reagents

The human gastric cancer cell line AGS was purchased from the American Type Culture Collection (ATCC, Manassas, VA, USA). GA0518 and GA0804 are patient-derived cells isolated from ascites or patient-derived xenograft (PDX) tumors from pancreatic cancer (PC) specimens and were described previously^33^. KP-Luc2 murine GAC cell line was from Dr. Jo Ishizawa in the Department of Leukemia of MDACC and was previously reported^34^. All cell lines were authenticated and profiled biannually by the Cell Line Core Facility at The University of Texas MD Anderson Cancer Center.

YAP antisense oligonucleotides (ASOs) and TAZ ASOs were sourced from Ionis Pharmaceuticals, Inc. (Carlsbad, CA, USA). The ASOs were initially dissolved in dimethyl sulfoxide (DMSO) at a concentration of 10 mM, with aliquots stored at −20°C. Fresh working solutions were prepared as required for each experiment.

### Isolation of GCPM cells from malignant ascites of GC patients

Malignant ascites (100 ml to 2000 ml) was collected under an IRB approved protocol (Lab01-543) at MDACC and CCCB0001 of the Cooper-Coriell Cancer Biobank at CCRC respectively. Cytology was confirmed by clinical pathologists (R.W&D.C). Ascites were centrifuged at 2000 rpm 20 min and and centrifuged PC cells and supernatant are processed and stored. Cell pellets are re-suspended in RPMI1640 + 5% FBS + 1% Penn/Strep and red blood cells (RBCs) are lysed in RBC lysis buffer. GCPM cell pellets are fixed and made to cell block and sectioned for evaluation of markers by IHC and Co-IF. Alive cells with 10% DMSO were stored at Liquid Nitrogen for planed flowcytometry and scRNA-seq later.

### Lentiviral Transfection and Establishment of Stable Cell Lines

YAP1 knockout cells were generated from the GA0518 cell line using the LentiCRISPR/Cas9 system. Briefly, gRNAs targeting YAP1 were designed using the MIT CRISPR Design Tool (http://crispr.mit.edu/). These gRNAs were cloned into the pLentiCRISPR v1 vector (GeCKO LentiCRISPR resource, MIT; http://genome-engineering.org/gecko/) using BsmBI digestion. The vector backbone, containing the Cas9 gene, was ligated with gRNA duplexes, and successful clones were verified by sequencing. Lentivirus was produced in HEK293T cells cultured in six-well plates by co-transfecting the pLentiCRISPR-gRNA construct with packaging plasmids pCMV.Dr8.2 and pCMV.VSV.G at a ratio of 10:10:1.

Lentiviral supernatant was used to transduce GA0518 cells expressing mCherry-Luciferase in six-well plates, supplemented with 8 µg/ml polybrene. Transduced cells were selected with puromycin at appropriate concentrations for 1-2 weeks to establish stable YAP1 knockout lines. Cells were subsequently propagated, and knockout efficiency was confirmed by Western blot analysis.

### Co-immunoprecipitation Assay

Co-immunoprecipitation (Co-IP) assays were performed using the Thermo Scientific™ Pierce™ Classic Magnetic IP/Co-IP Kit (#88804) according to the manufacturer’s instructions. Briefly, prepared cell lysates were incubated overnight at 4°C with antibodies against: TAZ (Novus, NBP1-85067), YAP1 (Cell Signaling Technology, #14074), TEAD1 (Cell Signaling Technology, #12292), TEAD2 (MilliporeSigma, Cat#SAB4503373), TEAD3 (Abcam, #ab75192), TEAD4 (Abcam, #58310), c-JUN (Cell Signaling Technology, #9615), or Fos-B (Cell Signaling Technology, #2251). Protein A/G magnetic beads were then added to the lysate-antibody mixtures and incubated for 1 h at room temperature to capture immune complexes. Beads were washed twice with IP Lysis/Wash Buffer and once with purified water to remove unbound proteins. Finally, antigen-antibody complexes were eluted from the beads for downstream analysis. Antibody concentrations used for subsequent Western blotting matched those employed in standard Western blot assays.

### Western Blotting

Proteins were extracted from whole-cell lysates using RIPA buffer. Protein concentration was quantified using a BCA Protein Assay Kit (Thermo Fisher Scientific) according to the manufacturer’s instructions, with bovine serum albumin (BSA) as the standard. Equal amounts of protein were separated by 10% polyacrylamide gel electrophoresis (SDS-PAGE) and transferred to PVDF membranes using a Trans-Blot® Turbo™ Transfer System (Bio-Rad). Membranes were incubated with the designated primary antibodies overnight at 4°C, followed by incubation with appropriate horseradish peroxidase (HRP)-conjugated secondary antibodies. Protein bands were visualized using ECL Western Blotting Detection Reagent (Thermo Fisher Scientific) via chemiluminescence. The following primary antibodies and dilutions were used: YAP1 (1:1000), TAZ (1:1000), TEAD1 (1:1000), TEAD2 (1:1000), TEAD3 (1:1000), TEAD4 (1:1000), Survivin (1:1000), Snail (1:500), β-actin (1:10,000).

### YAP/TEAD Luciferase Reporter Assay

YAP/TEAD transcriptional activity was assessed using the Dual-Luciferase Reporter Assay System (Promega). AGS, GA051816, and GA080417 cells were transiently co-transfected with UAS-luciferase and Gal4-TEAD plasmids, along with a Renilla luciferase vector as an internal control. For specific experimental conditions, a human YAP1 overexpression vector was also co-transfected. Eight hours post-transfection, cells were treated with various concentrations of antisense oligonucleotides (ASOs) or vehicle control for 24 h. Luciferase activity was then measured using a luciferase assay kit (Promega). Firefly luciferase activity was normalized to Renilla luciferase activity to account for transfection efficiency variations. Transfection experiments were independently performed at least three times, with each condition assayed in triplicate.

### Immunofluorescent Staining

Immunofluorescence staining was performed as previously described^5^. Antigen retrieval was conducted using Antigen Unmasking Solution (BioGenex Laboratories). The following primary antibodies were applied at the indicated dilutions: YAP1 (Cell Signaling Technology, #14074, 1:100), TAZ (Novus, NBP1-85067; 1:100); Ki67 (Thermo Fisher Scientific, #RM-9106-S1, 1:150), TEAD1 (Cell Signaling Technology, #12292, 1:100), TEAD2 (SAB4503373; 1:100), TEAD3 (Abcam, ab75192; 1:100), TEAD4 (Abcam, #58310, 1:100), CD8 (Cell Signaling Technology, #98941, 1:100), CD206 (Abcam, #64693, 1:100). Slides were mounted using Vectashield Mounting Medium with DAPI (Vector Laboratories) and visualized under a Nikon A1 confocal laser scanning microscope.

### Colony Formation Assay

GA051816 gastric cancer (GC) cells were seeded in six-well plates at a density of 800 cells per well (optimized to yield countable colonies). Cells were treated with YAP ASO, TAZ ASO, or a combination of YAP ASO + TAZ ASO at indicated concentrations. After 10–14 days of culture, colonies were fixed with 3% crystal violet in 10% formalin. Colonies containing >50 cells were counted, and clonogenic survival fraction was calculated. All treatments were performed in at least triplicate.

### Transwell Migration Assay

Migration assays were performed using 24-well Transwell® plates with 8.0-µm pore inserts (Corning). The lower chamber contained 750 µL of RPMI-1640 medium supplemented with 20% FBS. Target cells (1 × 10⁵) in RPMI-1640 medium with 1% FBS were seeded in the upper chamber. YAP ASO or TAZ ASO was added at specified concentrations. Plates were incubated at 37°C for 24 h. Non-migrated cells on the upper surface were removed by scraping. Migrated cells on the filter’s lower surface were fixed with 10% formalin and stained with 0.5% crystal violet. Images from the transwell migration assay were captured and analyzed with the’Analyze Particles’function in ImageJ software (NIH, Bethesda, Mary-land, USA), where colony counts per well were calculated automatically. Migrated cells on the filter’s lower side were quantified by counting five randomly selected fields under a microscope at 20×magnification. Each assay was performed in triplicate.

### RT-qPCR

Total RNA was extracted from cells or tissues using TRIzol™ reagent (Thermo Fisher Scientific). RNA concentration and purity were assessed by spectrophotometry (NanoDrop). Reverse transcription was performed using the LunaScript® RT SuperMix Kit (New England Biolabs) according to the manufacturer’s protocol. RT-qPCR was performed on the cDNA using SYBR Green Master Mix (Applied Biosystems). Gene expression levels were normalized to GAPDH expression using the 2-△△Ct method. Each assay was conducted in triplicate. Primer sequences are shown in Supplemental Table 1.

### Single Cell RNA Sequencing (sc-RNA Seq) Analysis

Droplet-based 3’single-cell RNA-Seq (10x Genomics) was performed on ascites samples (n = 20) at SMF core at MD Anderson Cancer Center as described^35^. In brief, transcripts were mapped, assigned to individual cells by barcodes using Cell Ranger, and analyzed using Seurat in R 3.5.0. Genes with detected expression in at least 3 cells, and cells with at least 200 genes detected were used. The first 15 Principal Components were used for clustering (resolution = 0.6), RNA Sequencing analysis Total RNA was isolated using a miRNeasy Mini Kit (Qiagen) according to the manufacturer’s protocol from all ascites cells. Only RNA with more than 7 of RNA integrity number was sent to DNA core for the RNA Sequencing. The raw RNA-sequencing (RNA-seq) readouts were mapped to the human GRCh38 assembly reference genome using TopHat2, an open-source software tool that aligns RNA-seq reads to a reference genome. The heatmaps were generated with unsupervised clustering and the differential gene expression analyses were performed with DESeq2 (R/Bioconductor package) using adjusted p value < 0.05 as the significance cutoff.

### Immunohistochemistry

IHC for YAP1 or TAZ was performed on human tissue microarray slides consisting of samples of around 390 gastric tumor tissues and non-neoplastic gastric tissues from patients underwent total or subtotal gastrectomy for GC between January 2006 and December 2008 at the First Affiliate Hospital of China Medical University. Wrtieen informed consents were obtained from all patients and all patients were followed up via telephone inquiry or questionnaires. Antibodies for YAP1 was described previously^5^. Sections were incubated with primary antibodies: human YAP1 antibody (Cell signaling, cat#14074; 1:100) and human TAZ antibody (Novus, cat#NBP1-85067; 1:100) followed by biotinylated secondary antibodies, and streptavidin HRP kit (Vector Laboratories, PK-6100). IHC score was based on the criteria based on our previous study^5,36,37^.

### The PDX and KP-Luc2 syngeneic mouse models

All animal experiments were approved by the Institutional Animal Care and Use Committee (IACUC) and conducted in compliance with relevant ethical guidelines. Ten- to twelve-week-old SCID mice were subcutaneously inoculated with GA051816 cells (2 × 10⁶ cells/mouse; n = 5 per group). For the YAP/TAZ ASO efficacy study, treatment commenced 15 days post-inoculation when xenografts were established. Mice received intraperitoneal injections of: YAP ASO (40 mg/kg) TAZ ASO (40 mg/kg). Combination therapy (YAP ASO 20 mg/kg + TAZ ASO 20 mg/kg); PBS vehicle control (100 μL/mouse). Treatments were administered five times weekly for three consecutive weeks.

For the combination therapy in KP-Luc2 syngeneic model, established KP-Luc2 xenograft-bearing mice were treated with: YAP ASO monotherapy (40 mg/kg), anti-PD-1 neutralizing antibody (Bio X Cell; Cat# BE0146-R-100mg; 10 mg/kg, IP), Combination therapy (YAP ASO 40 mg/kg + anti-PD-1 10mg/kg) and PBS control group 5 times a week for total 3 weeks. Tumor volumes (calculated as length × width² × 0.5), tumor weights, and body weights were monitored throughout the study period as previously described^38^.

### Statistical Analyses

All statistical analyses were conducted using SPSS software (version 20.0; IBM Corp.). Continuous data are presented as mean ± standard deviation (SD) or standard error of the mean (SEM), as specified in the figure legends. For comparisons between two independent groups, we employed unpaired two-tailed Student’s t-tests after verifying the assumptions of normality and homogeneity of variance. The threshold for statistical significance was established a priori at p < 0.05 for all analyses. The specific sample size for each experimental condition is provided in the respective figure legends. All statistical tests were selected based on the experimental design and data characteristics, with appropriate validation of test assumptions.

## Supplemental Figure Legends

**Supplemental Figure 1.**
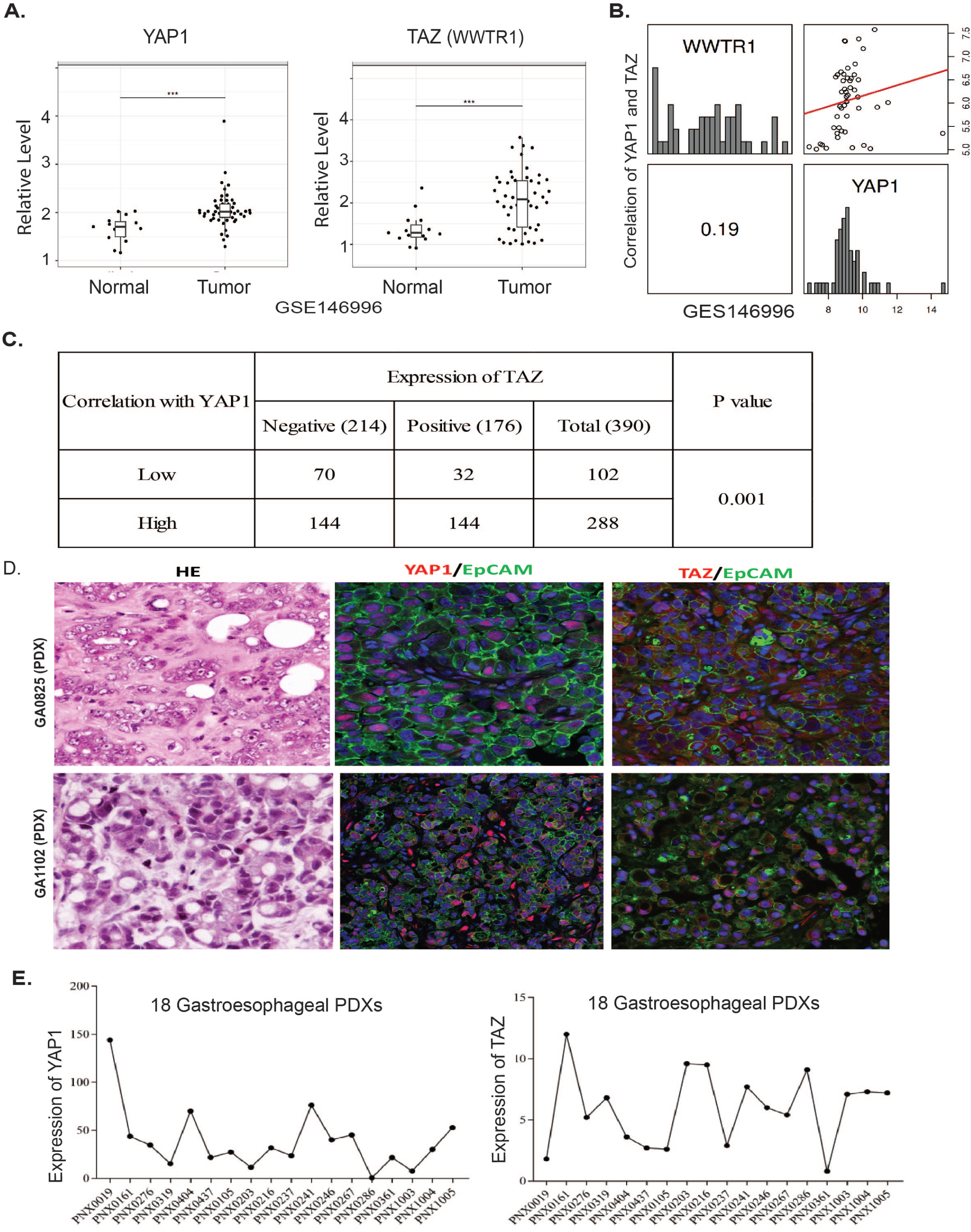
YAP1 and TAZ are highly expressed in GC tumor tissues and PDXs. A. Increased expression of YAP1 or TAZ (WWTR1) in GC tumor tissues compared to normal tissues from a public GC data set (GSE146996). B. Both YAP1 and TAZ are correlated (R=0.19) in the same dataset. C. Expression of YAP1 and TAZ in our own TMA of 390 GC Cases TAZ was statistically analyzed (p=0.001); D. co-expression of YAP1/EpCAM or TAZ/EpCAM was determined by co-IF in two representative PDXs; E. mRNA expression of YAP1 or TAZ was determined in tumor tissues of 18 PDXs from gastroesophageal cancers.

**Supplemental Figure 2.**
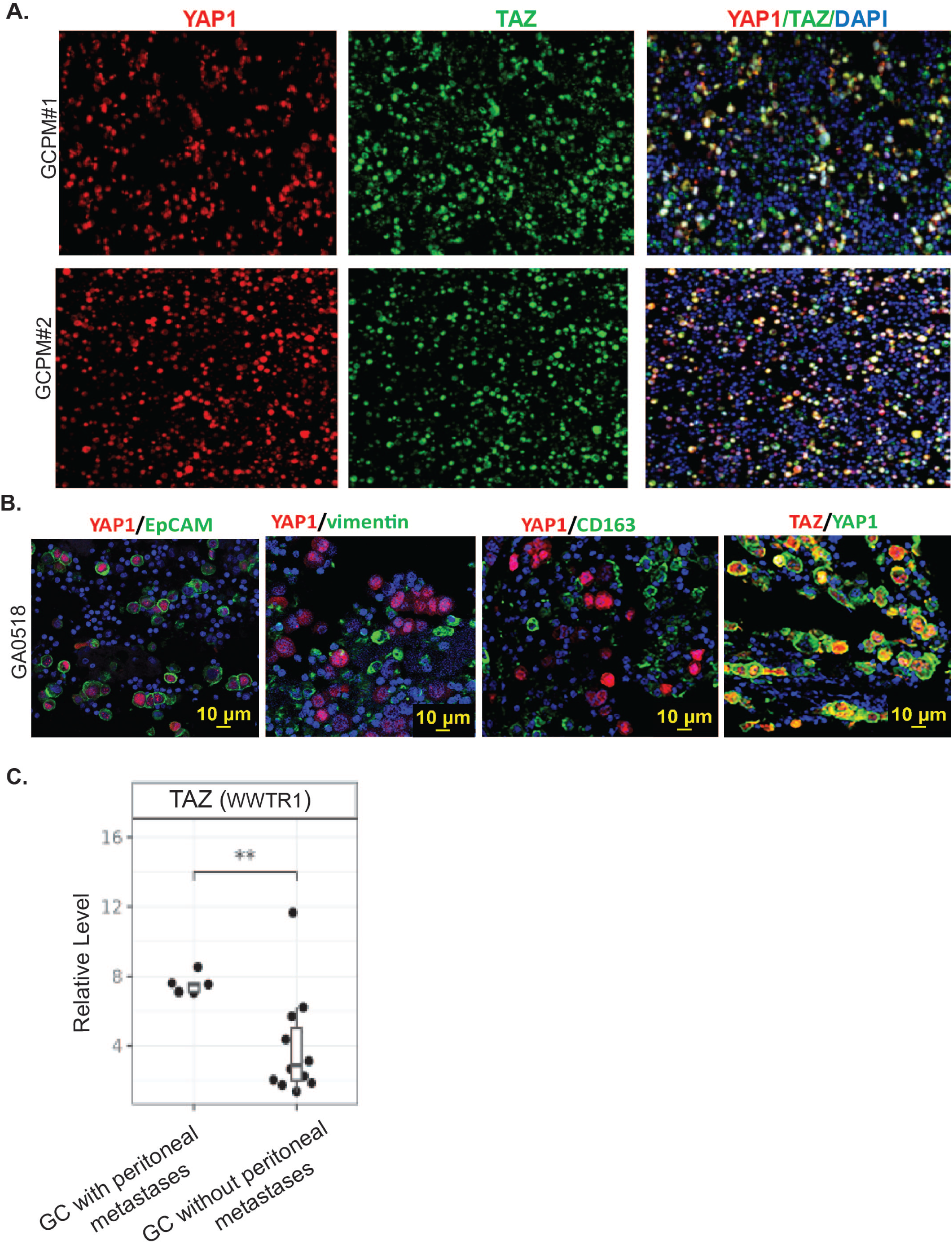
Co-expression of YAP1 and TAZ in representative GCPM samples. A. coexpression of YAP1 or TAZ in nuclear of two representative GCPM specimen was determined by co-IF staining; B. Coexpression of YAP1 with epithelial tumor marker EpCAM or TAZ or stromal marker vimentin and M2 macrophage marker CD163 in GA0518 GCPM cases; C. TAZ expression was determined in GC with peritoneal metastases compared to GC without peritoneal metastases from a public GC dataset GSE289037.

**Supplemental Figure 3.**
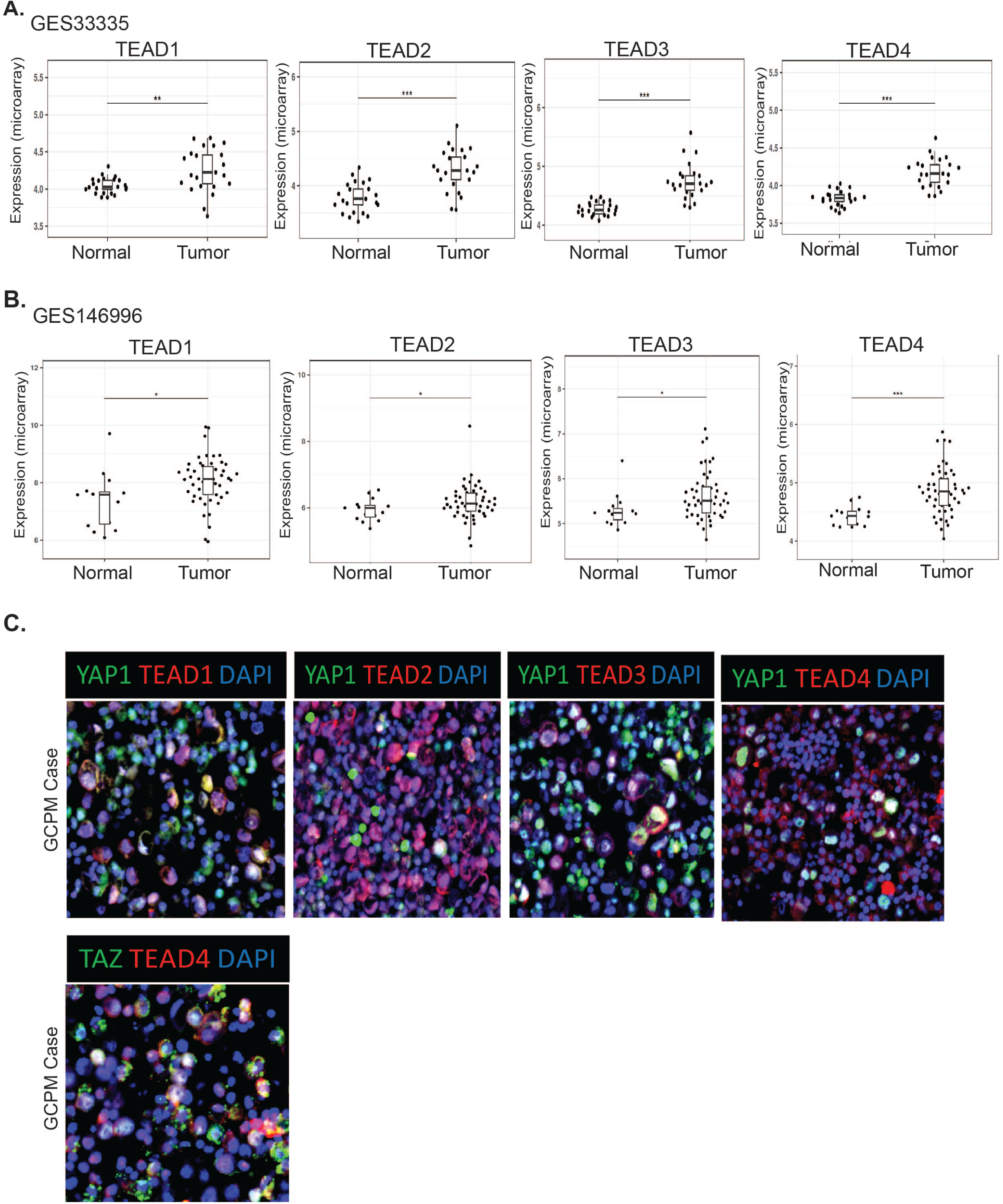
High expression of TEAD1-TEAD4, the transcription factors (TFs) for YAP1/TAZ in primary GC and GCPM cases. A. Expression of TEAD1, TEAD2, TEAD3 and TEAD4 was significantly higher in GC tumor tissues compared to normal by analyzing GES33335 GC dataset; B. Expression of TEAD1, TEAD2, TEAD3 and TEAD4 was significantly higher in GC tumor tissues compared to normal by analyzing another GC cohort GES146996; C. Co-IF staining of YAP1 and TEAD1-4 or TAZ with TEAD4 in representative GCPM cases.

**Supplemental Figure 4.**
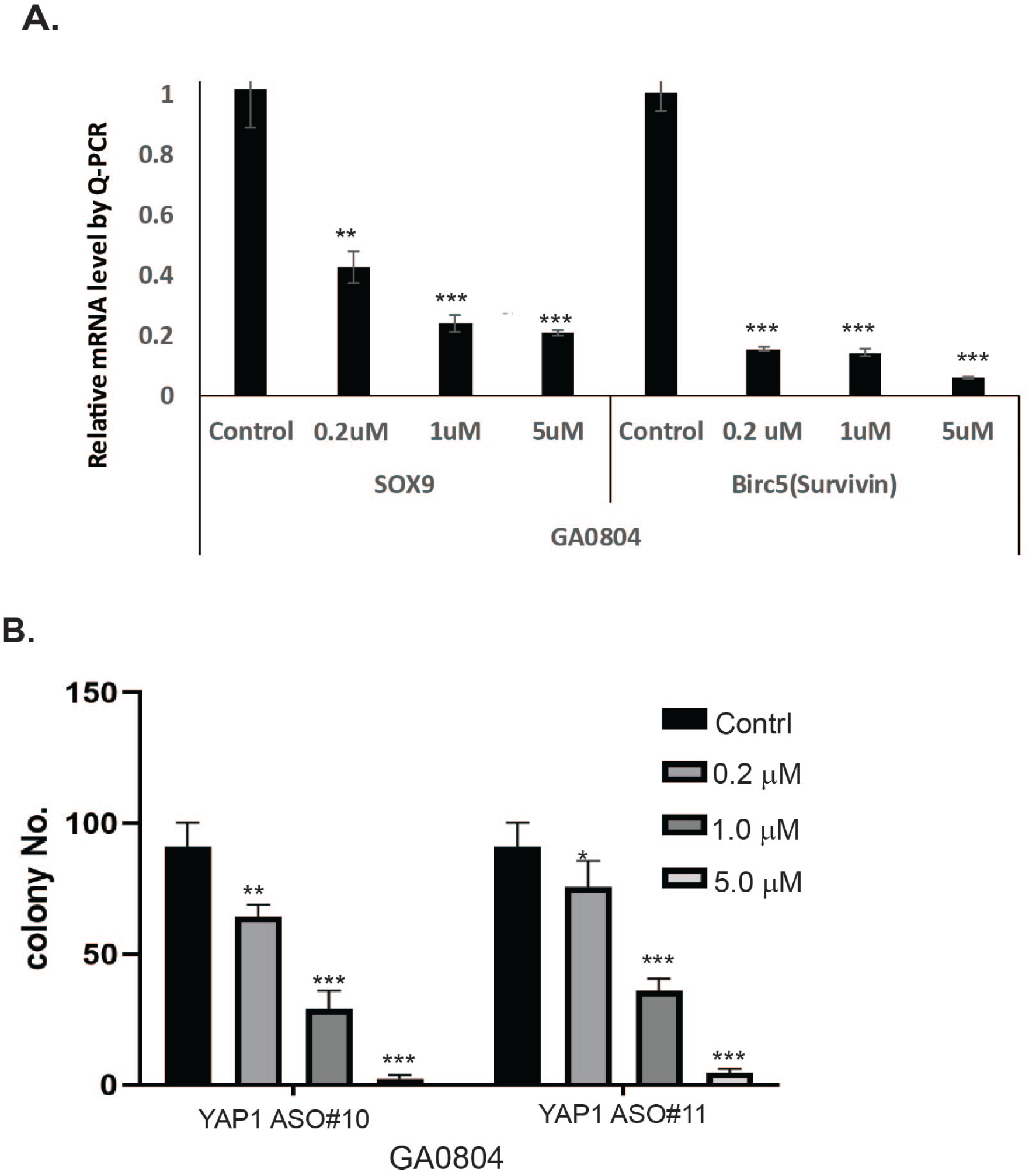
Effects of YAP1 ASO on YAP1 targets and on tumor cell colony formation. A. mRNA levels of SOX9 and Birc5 were dose-dependently decreased by YAP1 ASO as determined by q-PCR in GA0804 cells **p<0.01; ***p<0.001; B. Colony formation was determined in GA0804 cells upon treatment of YAP1 ASO#10 or YAP1 ASO#11 at different dosage as indicated. **p<0.01; ***p<0.001.

**Supplemental Figure 5.**
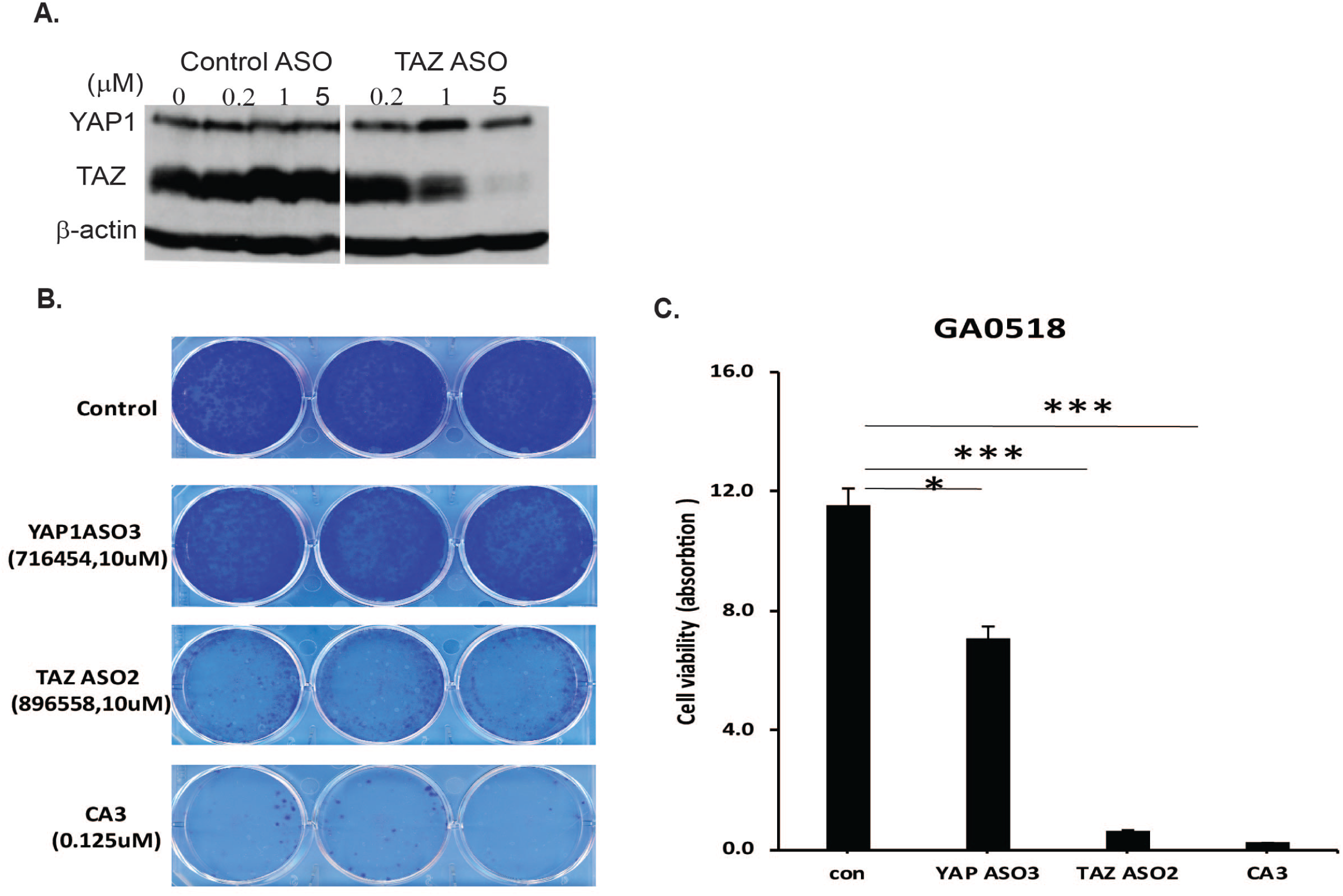
TAZ ASO specifically suppress TAZ expression and inhibit GA0518 cell colony formation. A. Expression of YAP1 and TAZ was determined in GA0518 cells by western blot after treatment of TAZ ASO in dosage as indicated; B&C. Demonstration of colony formation (B) and quantification (C) in GA0518 cells treated with YAP1 ASO, TAZ ASO or a reported YAP1/TEAD inhibitor CA3 at the dosage indicated; *p<0.05, **p<0.01; ***p<0.001.

**Supplemental Figure 6.**
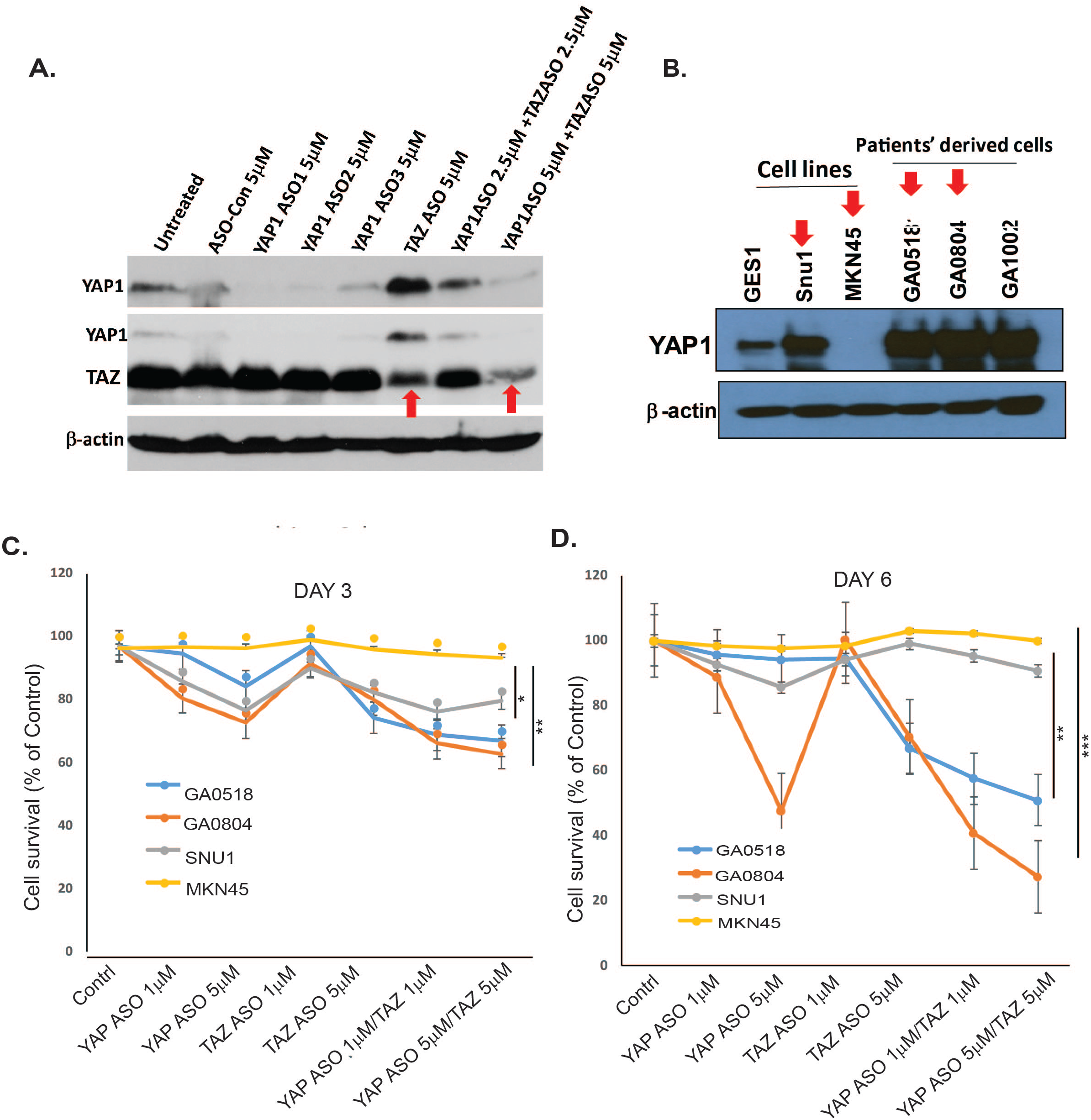
Cotreatment GC tumor cells using YAP1, and TAZ ASOs effectively suppress both YAP1 and TAZ expression and suppress tumor cell growth in YAP1 high GC cells. A. Expression of YAP1 and TAZ was determined in GA0518 cells by western blot after treatment of three YAP1 ASOs, TAZ ASO and their combination at the dosage as indicated; **B**. Expression of YAP1 in GES1, two GC cell lines and three GCPM derived tumor cells was determined by Western blot; **C&D**. Cell survival was detected by MTS assay in four GC cell lines with different YAP1 levels at dosage indicated. *p<0.05; **p<0.01; ***p<0.001.

**Supplemental Figure 7.**
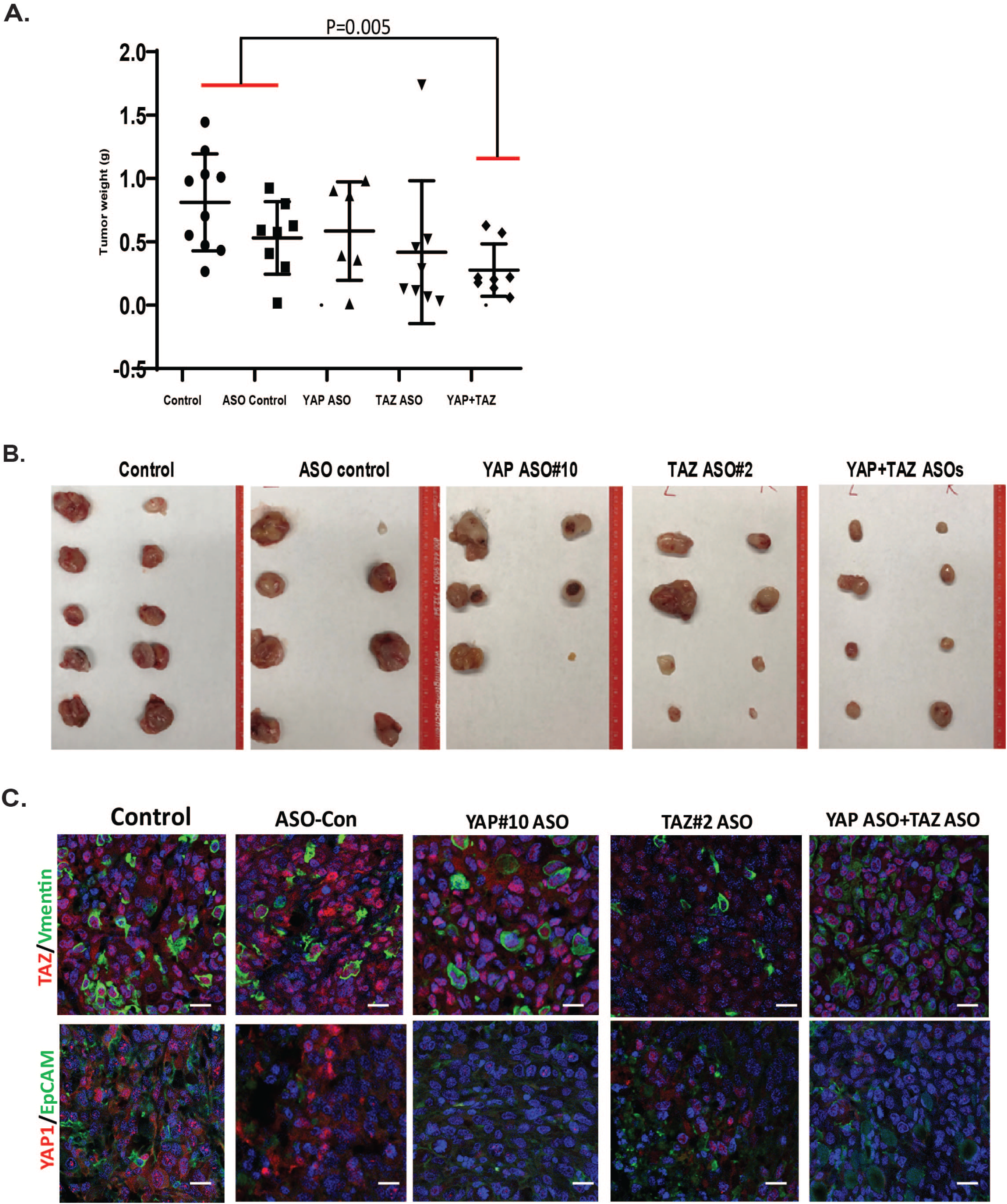
ASO co-targeting of YAP and TAZ significantly attenuated tumor growth in additional PDX. A. Combined ASO inhibition of YAP1 and TAZ suppressed tumor weights in a GC PDX model; YAP1 ASO or TAZ ASO: 50mg/kg, the combination of YAP1 ASO (25mg/kg) and TAZ ASO (25mg/kg); 5 times a week for three weeks. B. Actually tumor sizes in different treatment groups were shown at the end of the experiment. C. co-expression of TAZ/vimentin and YAP1/EpCAM as determined by IF staining in YAP1 ASO, TAZ ASO, or combination ASO-treated PDX tumors; Scale bar: 20 μm;

---

**Supplemental Table 1.**
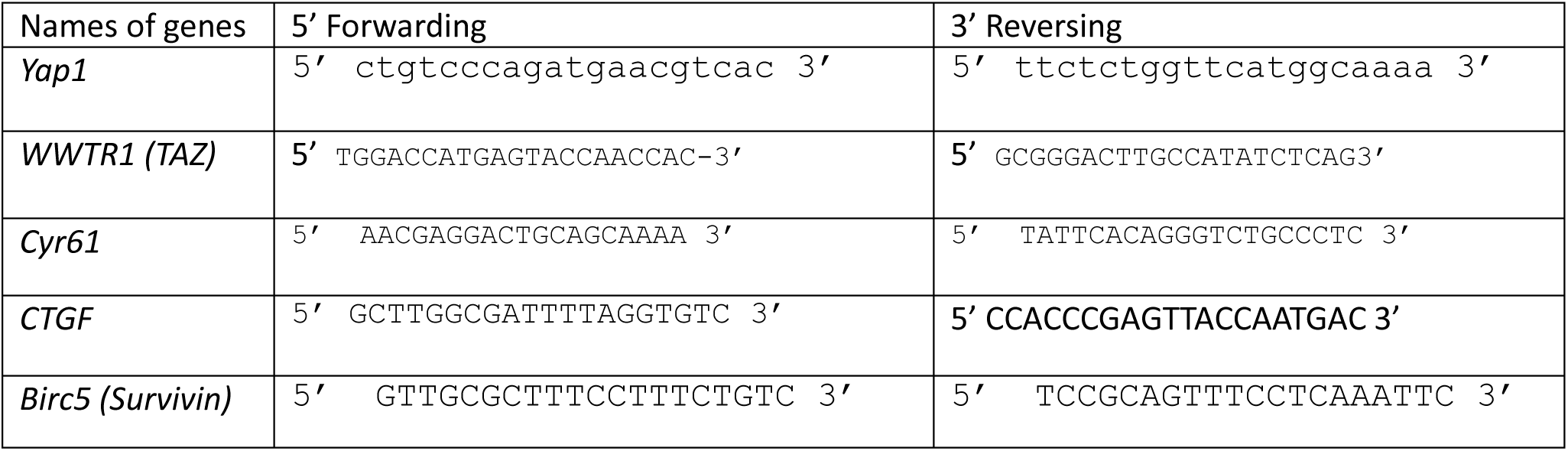
Primers used in this study.

**Supplemental Table 2.** Antibodies used in this study for Western blot, Immunoprecipitation, and Immunofluorescent staining.

| Antibody | Catalog No. | Company Name |
| --- | --- | --- |
| YAP1 | #14074 | Cell Signaling |
| TAZ | NBP1-85067 | Novus |
| TEAD1 | #12292 | Cell Signaling |
| TEAD2 | SAB4503373 | Millipore Sigma |
| TEAD3 | ab75192 | Abcam |
| TEAD4 | ab58310 | Abcam |
| c-Jun | #9615 | Cell Signaling |
| FOSB | #2251 | Cell Signaling |
| Ki-67 | RM-9106-S1 | Thermo Scientific |
| CD8 | #98941 | Cell Signaling |
| CD163 | #25121 | Cell Signaling |
| CD206 | ab64693 | Abcam |
| Survivin | #2808 | Cell Signaling |
| SOX9 | AB5535 | Millipore Sigma |
| EpCAM | #2929 | Cell Signaling |
| Vimentin | sc-66002 | Santa Cruz Biotechnology |
| Snail | sc-271977 | Santa Cruz Biotechnology |
| β-Actin | MAB8929 | Biotechne, R&D Systems |

